# Population-level control of two manganese oxidases expands the niche for bacterial manganese biomineralization

**DOI:** 10.1101/2024.09.19.613919

**Authors:** Gaitan Gehin, Nicolas Carraro, Jan Roelof van der Meer, Jasquelin Peña

## Abstract

The enzymatic oxidation of aqueous divalent manganese (Mn) is a widespread microbial trait that produces reactive Mn(III, IV) oxide minerals. These biominerals drive carbon, nutrient, and trace metal cycles, thus playing important environmental and ecological roles. However, the regulatory mechanisms and physiological functions of Mn biomineralization are unknown. This challenge arises from the common occurrence of multiple Mn oxidases within the same organism and the use of Mn oxides as indicators of combined gene activity. Through detection of gene activation in individual cells, we discover that expression of *mnxG* and *mcoA*, two Mn oxidase-encoding genes in *Pseudomonas putida* GB-1, is confined to subsets of cells within the population, with each gene showing distinct spatiotemporal patterns that reflect local microenvironments. These coordinated intra-population dynamics control Mn biomineralization and illuminate the strategies used by microbial communities to dictate the extent, location and timing of biogeochemical transformations.

## Introduction

Life and minerals are inextricably linked^1–8^. In microorganisms, biomineralization—the process of mineral precipitation through biological activity—serves many physical, chemical, and biological functions^2^. These functions include protection from UV radiation, antibiotics, or encrustation; detoxification of reactive oxygen species, CO2 or other toxic compounds; and storage of nutrients, carbon or energy. Biomineralization also supports microbial metabolism by allowing some microorganisms to generate additional proton motive force, conduct extracellular electron transfer, or colonize environmental niches at redox interfaces^2,9–13^. For some systems such as manganese (Mn)^14^, however, the mechanisms and functions of biomineralization are partially or scarcely known.

Manganese biomineralization is a widespread microbial trait performed by a phylogenetically diverse group of bacteria and fungi^15–18^. Mn-oxidizing microorganisms are found in numerous aquatic and terrestrial environments^5,19,20^. They typically use one or more multicopper oxidase (MCO) or animal heme peroxidase^15,18^ enzymes to oxidize aqueous Mn(II) to Mn(III, IV) species, which subsequently precipitate as reactive layer-type Mn oxides^5,20^. A recent genome meta-analysis showed the wide conservation of multicopper Mn-oxidase genes in 682 out of 2197 tested bacterial genomes, and found co-occurring multicopper oxidases in a third of the predicted Mn oxidizers^15^. The enzymatic reaction involves extracellular or membrane-bound enzymes and electron transfer chains that donate up to two electrons from Mn(II) species to molecular oxygen and, in some cases, H2O2^17^. The rates of enzymatic Mn(II) oxidation are up to five orders of magnitude faster than expected from homogeneous aqueous-phase reactions or heterogeneous reactions between aqueous and solid-phase species^21^, suggesting that this process has one or more critical biological functions. However, how and under which extracellular environmental conditions the expression of Mn oxidases is regulated by the cell, how this contributes to mineral precipitation, and whether this process benefits microbial life remains largely unknown.

A key challenge in advancing our understanding of the functional role(s) of bacterial Mn biomineralization is that this complex process cannot be measured or described by any single endpoint. Mn-oxidizing enzymes have been purified or partially purified, and their catalytic activity has been studied, notably with model Mn oxidizers such as *Bacillus sp.* PL-12^14,22,23^, *Roseobacter* AzwK-3b^24,25^, and especially *Pseudomonas putida* GB-1^15,26–32^. Nonetheless, to address the functional and ecological roles of Mn oxidation and biomineralization, the enzymes need to be studied at the individual cell and population level. To date, most studies of enzymatic manganese oxidation and biomineralization rely on the quantification of the Mn oxide itself, which is problematic for a number of reasons. Biogenic Mn oxides form extracellularly^17,22,26,29,33^ and often in a matrix of extracellular polymeric substances (EPS), which results in complex microbe-mineral assemblages^34^. Manganese oxides are also highly reactive towards metals^35,36^ and are susceptible to reductive dissolution in the presence of organic compounds and extracellular metabolites such as sugars, organic acids, and siderophores^17,37–39^. Therefore, using redox-sensitive Mn oxides as indicators of Mn oxidase gene expression and enzymatic activity can lead to an incomplete understanding of the mechanisms of Mn oxidation. This approach is further obfuscated by the presence of multiple Mn oxidases and multiple regulation pathways within the same organism^15,24,27,40^. To overcome these challenges, two advanced keys are required: the enzymatic activity must be decoupled from Mn oxide formation, and Mn-oxidase gene activation must be linked to mineral precipitation at the level of individual cells within bacterial populations.

Our goal here was thus to develop a system to disentangle the dynamics of Mn-oxidase gene activation alongside Mn oxide formation, as a function of environmental growth conditions. As a model system, we used *Pseudomonas putida* GB-1, a bacterium that has been shown to possess three genes for Mn oxidation: two multicopper oxidases (MnxG and McoA) and a heme peroxidase (MopA)^28^. Both MnxG and McoA can catalyze Mn oxidation independently, as shown in a study of *P. putida* GB-1 derivatives with in-frame deletions of either *mnxG* or *mcoA*^26^. To follow Mn-oxidase gene activation at the single cell level, we constructed reporter gene fusions in wild-type *P. putida* GB-1 containing the full Mn-oxidizing machinery, consisting of a single copy, a chromosomally-integrated fusion of the promoter upstream of either *mnxG* or *mcoA* (i.e., *P2447_mnxG* and *P2665_mcoA*, hereafter *PmnxG* and *PmcoA*). We did not select *mopA* because it has never shown any activity in the wild-type strain or mutants lacking both *mnxG* and *mcoA*^42^. We anticipated that cells activating either promoter would trigger the formation of the reporter fluorescence, which serves as a proxy for Mn-oxidase expression. Fluorescent reporters also provide signals that can be quantified using microscopy in real-time in individual cells, allowing us to detect Mn-oxidase activation over time as a function of spatial position. To create different physiological conditions, we cultured GB-1 on surfaces and in liquid suspension to follow Mn-oxidase gene expression in microcolonies as well as in individual planktonic cells and cell aggregates (*P. putida* GB-1 quickly forms strongly adhering multi-cell aggregates^34^), respectively.

We show that *mnxG* and *mcoA* are successively activated in non-dividing cells in stationary phase, only in the presence of Mn, and independently of the growth condition. By simultaneously localizing Mn oxide formation to reporter expression, we find that MnxG is responsible for the initial precipitation of Mn oxides, and McoA contributes to biomineralization under conditions where MnxG activity is restricted. Mn-oxidase gene activation and mineral precipitation occurred only in a subpopulation of cells, whose proportion is not dependent on planktonic or sessile lifestyle, but rather is determined by local environmental conditions. The discovery of subpopulation-dependent Mn-oxidase expression in GB-1 provides a new framework for understanding the cellular function(s) of Mn oxidation and biomineralization, which we hypothesize may involve cellular cooperation to dictate the need, location, and timing of the Mn transformation reactions.

## Results

### Decoupling gene activation from Mn oxide precipitation

To facilitate the detection of Mn oxidase gene expression in *P. putida* GB-1, we constructed two derivative strains with eCherry fused to the isolated promoters of either *mnxG* or *mcoA* (**Supplementary** Figs. 1 and 2). Reporter fusions were placed in a single gene copy on the *P. putida* GB-1 chromosome and shielded for upstream and downstream transcription readthrough. These derivative strains contain the complete Mn-oxidizing machinery and report either *mnxG* or *mcoA* activation (hereafter, bioreporters). Sequence analysis revealed a similar structure for the two promoter regions upstream of *mnxG* (*PmnxG*) and *mcoA* (*PmcoA*), with each containing the predicted binding sites for integration host factor (IHF) and sigma-54 (σ^54^) transcription factor (**Supplementary** Fig. 1). The binding site suggested that both genes are IHF-σ^54^ dependent promoters, whose activation is frequently associated to stationary phase conditions^43,44^ and regulated by specific environmental signals^45–48^. Reporter gene activation in both strains then serves as a proxy for Mn-oxidase gene expression that can be compared to the precipitation of Mn oxides, allowing us to quantify biological and geochemical expressions of Mn biomineralization.

### Mn triggers the activation of *PmnxG* and *PmcoA* promoters

To examine the activation of *mnxG* and *mcoA* promoters, we recorded eCherry fluorescence signals in cells of wild-type *P. putida* GB-1, the *PmnxG* bioreporter, and the *PmcoA* bioreporter after 48 h in the presence or absence of 50 µM MnCl2. Experiments were carried out using cells grown on solid agarose surfaces (**Fig. 1**) or in liquid-suspended culture (**Supplementary** Fig. 3). In the absence of Mn, there was no difference between the fluorescence intensity of the bioreporters and the wild type auto-fluorescence in the eCherry wavelengths (**Fig. 1a, b**). In the presence of 50 µM MnCl2, the median bioreporter signal for *PmnxG* in surface-grown microcolonies increased by 12.5- fold, and that of *PmcoA* increased by 4.7-fold relative to the no Mn condition (**Fig. 1a, b**). In liquid-suspended culture, the median fluorescence signal of the *PmnxG* bioreporter in the presence of 50 µM MnCl2 increased by 16-fold and that of *PmcoA* by 13-fold, compared to the no Mn condition (**Supplementary** Fig. 3). These results show that Mn is required for the activation of the *mnxG* and *mcoA* promoters.

**Fig. 1.**
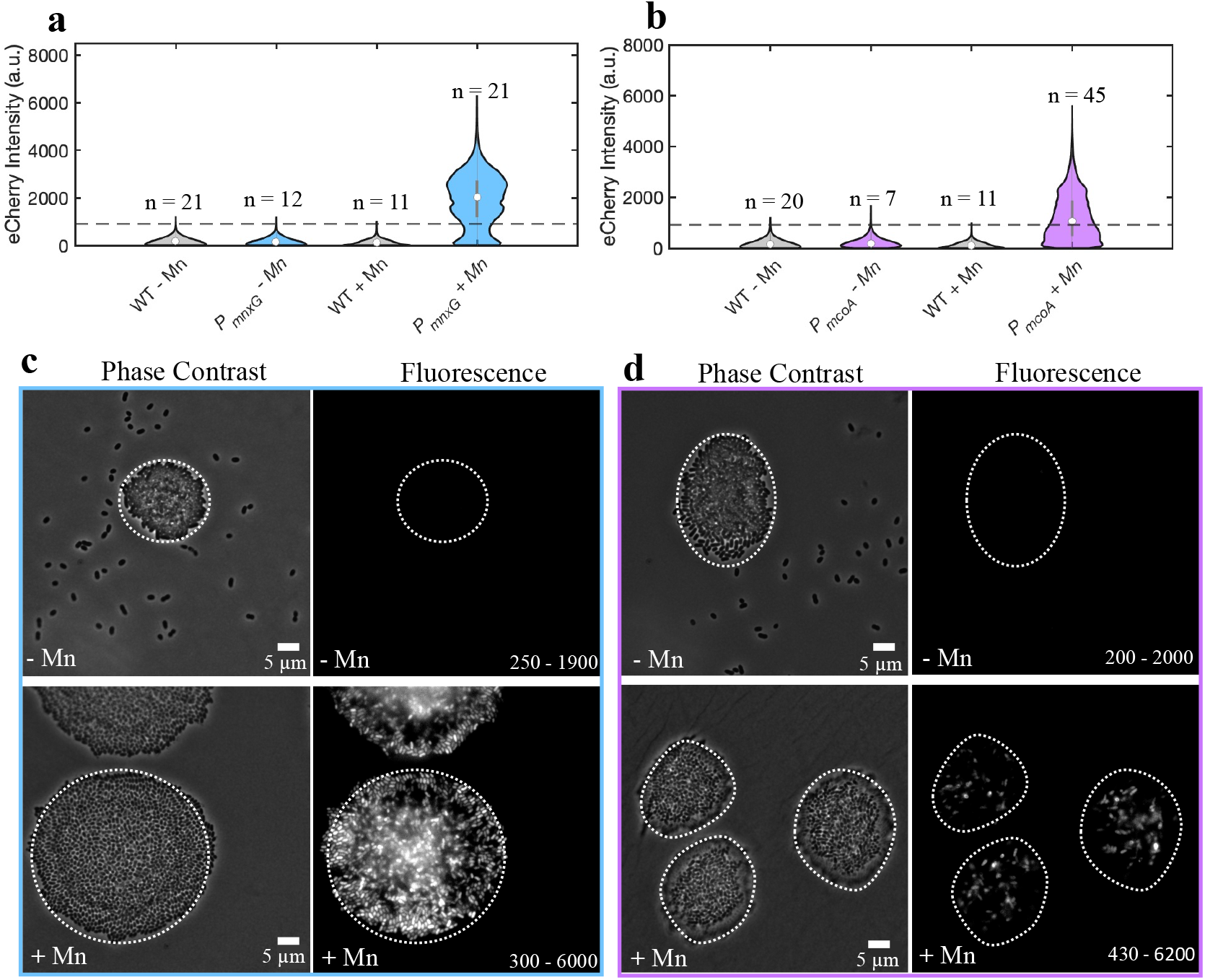
Expression of Mn oxidase gene promoters in the presence and absence of Mn(II) in microcolonies of *P. putida* GB-1. **a-b** Violin plot of the fluorescence intensity distribution in microcolonies of wild-type (WT) and bioreporter (*PmnxG*, **a**; *PmcoA*, **b**) strains without Mn and with 50 µM Mn(II), where n is the number of microcolonies analyzed for each condition. The eCherry fluorescent signal represents the pixel intensity distribution within the boundaries of the microcolonies in the phase contrast images. The fluorescence threshold is set as the 99^th^ quantile of the wild type distribution, calculated using background-subtracted images. Outliers removed at 6 × the *SD* of the respective distributions. **c-d** Representative phase contrast and fluorescence images of 48-h-grown *P. putida* GB-1 *PmnxG* (**c**) and *PmcoA* (**d**) microcolonies grown in the absence of Mn (top panels) and in the presence of 50 µM Mn(II) (bottom panels). Values on the fluorescence micrographs represent the range of lowest to highest pixel fluorescence signal intensity in the displayed image.

### Stationary phase and subpopulation-dependent activation of *PmnxG and PmcoA*

The bioreporter signals of individual cells within microcolonies grown in the presence of Mn, measured after 48 h, differed widely (**Fig. 1c, d**). Specifically, the distribution of pixel intensities in 21 microcolonies of the *PmnxG* bioreporter showed three distinct modes: one below the fluorescence threshold and two above the fluorescence threshold. This pattern in the fluorescence signal distribution within the microcolonies suggests bimodal gene activation, where *PmnxG* is inactive in a subpopulation of cells and active in another. The occurrence of two modes at 1600 a.u. and 2500 a.u. may result from the increased stacking of cells near the microcolony center relative to the microcolony edges. The pixel intensities of the *PmcoA* bioreporter in 45 microcolonies grown in the presence of Mn showed two modes, one at 60 a.u. and another at 2240 a.u.. The higher proportion of fluorescence intensities below the threshold than above the threshold suggests that the *PmcoA* promoter was inactive in most of the population and less transcribed overall than the *PmnxG* promoter (**Fig. 1a, b**).

To confirm the bimodality of the gene activation pattern for *PmnxG* and *PmcoA*, we monitored bioreporter activation over time in growing microcolonies. The maximum exponential growth rates (µ), calculated as the increase in projected surface area over time, were comparable for both bioreporters, with an average of µ = 0.19 ± 0.02 h^-1^ for strain *PmnxG* and µ = 0.13 ± 0.07 h^-1^ for strain *PmcoA*, and entry into stationary phase at 19.5 h and 19.0 h, respectively, for strain *PmnxG* and strain *PmcoA* (**Fig. 2a, b**). The timing of microcolony growth compared to the appearance of the reporter signals indicated that both promoters were activated exclusively after entry into stationary phase (**Fig. 2a, b**), as confirmed further by the disappearance of the fluorescence reporter signal in stationary phase cells exposed to fresh growth medium (**Supplementary** Fig. 4). However, the fluorescence signal from the *PmnxG* bioreporter started to appear 8 h after the microcolonies entered the stationary phase (**Fig. 2a**), whereas the signal from the *PmcoA* bioreporter only appeared 13 h after entry into stationary phase (**Fig. 2b**). In addition, the maximum fluorescence intensity was reached much earlier in the *PmnxG* bioreporter than the *PmcoA* bioreporter (**Supplementary** Fig. 5a, b), indicating that the timing and rates of expression for both Mn-oxidase genes must be different.

**Fig. 2.**
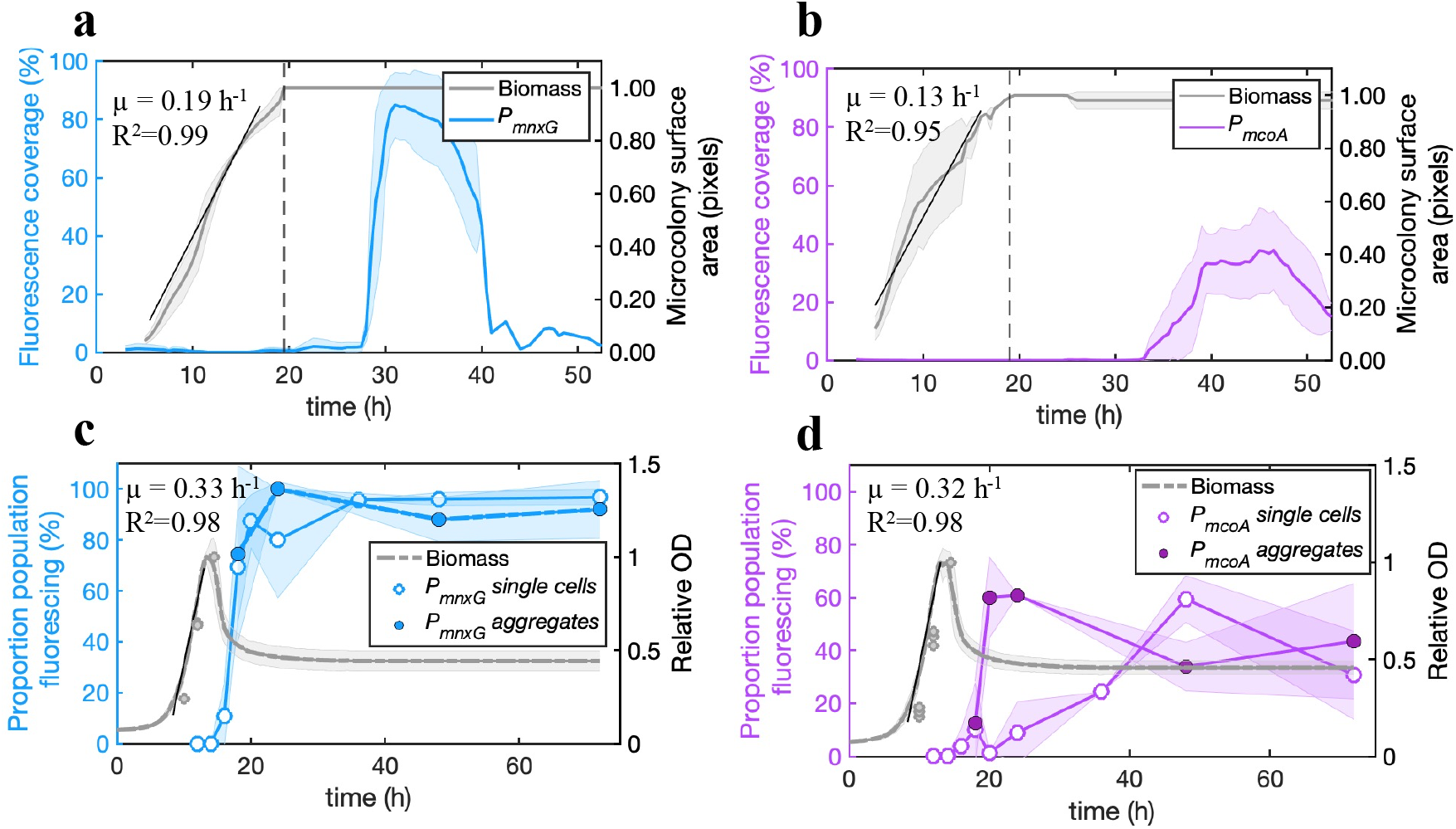
Temporal activation of Mn oxidase gene promoters of *P. putida* GB-1 in surface-grown microcolonies and liquid-suspended cultures. a-b Surface area of *PmnxG* (**a**) *PmcoA* (**b**) fluorescing cells (in blue or magenta) relative to the total microcolony surface area (in grey) at entry into stationary phase as indicated by the dotted lines, expressed as a percent (%). The growth curves, calculated as the increase in microcolony surface area over time, represent the average of 13 replicates for *PmnxG* and 21 replicates for *PmcoA*. Shaded areas represent the 95^th^ confidence interval. Growth rates (µ) are calculated during the exponential phase, as indicated by the black line and its linear regression coefficient (R^2^). **c-d** Proportion of individual (open symbols) or aggregate cells (closed symbols) fluorescing over time (blue and magenta lines) in liquid-suspended cultures for (**c**) *PmnxG* and (**d**) *PmcoA*. The shaded areas represent the standard deviation within triplicates. Relative OD corresponds to the culture turbidity (as OD600, in grey) normalized by the maximum OD600. Growth curves for liquid-suspended cultures were measured in 5 replicates using a plate reader. The decrease in relative OD after the exponential phase reflects cell aggregation. Separate relative OD600 measurements for cultures grown in flasks are shown by the gray symbols.

The total reporter fluorescence from the *PmnxG* microcolonies increased by 29-fold from the start of the activation to its maximum (**Supplementary** Fig. 5a), but the average fluorescence signal per individual reporting cell in this time interval was approximately constant (**Supplementary** Fig. 6a and c). Similarly, for the *PmcoA* bioreporter, the total fluorescence signal increased by 35-fold (**Supplementary** Fig. 5b), but the average signal per reporting cell varied by less than 5-fold (**Supplementary** Fig. 6b, d). After reaching saturation, the fluorescence signal in both cases decreased, indicating that promoter activity ceased or was exceeded by photobleaching due to fluorescence excitation (**Supplementary** Figs. 7 and 8). These results confirm that the increase in total fluorescence from both promoters was driven by an increase in the proportion of activated reporter cells rather than by the increase of fluorescence in individual reporting cells.

Based on the proportion of pixels with fluorescence values above the threshold value, we calculated the maximum proportion of reporting cells within stationary phase microcolonies at 85.1% for the *PmnxG* bioreporter and 37.6% for the *PmcoA* bioreporter (**Fig. 2a, b**). The 95^th^ confidence interval for the maximum active subpopulation was within 9.0% of the mean value for *PmnxG* (n = 13 microcolonies; **Fig. 2a**) and 15.1% of the mean value for *PmcoA* (n = 21 microcolonies; **Fig. 2b**). Image analysis of cells grown in liquid-suspended culture revealed similar proportions of cells activating either of the promoters (**Fig. 2c, d**). However, in liquid cultures at stationary phase conditions, nearly all cells activated the *PmnxG* promoter (**Fig. 2c**), whereas only about 60% activated the *PmcoA* promoter (**Fig. 2d**). Moreover, the rate of activation of the *PmcoA* promoter varied with the degree of cell aggregation, such that gene activation was faster within large aggregates (60% of the projected surface area after 20 h) than in planktonic cells (60% of the population activated after 48 h; **Fig. 2d**). To explore whether the absence of Mn-oxidase gene activation is associated to cell damage, we compared viability of cells with or without fluorescent reporter signal from the same culture. Cells grown in the presence of 50 µM Mn(II) for 48 h were seeded on agarose surfaces to follow cell division in real-time. These experiments showed no significant difference in the length of the lag phase or average division time between founder cells with or without previous fluorescent reporter signal, suggesting that reporter expression (and, by analogy, the respective Mn-oxidase gene activation) is not linked to cell viability (**Supplementary** Fig. 9). Therefore, the observed differences in the timing and proportion of cells showing Mn-oxidase promoter activation among individual cells in stationary phase is not due to differences in cell growth or viability but must reflect an underlying gradient or change in environmental conditions experienced by the cells.

### Spatiotemporal patterns in Mn oxidase gene expression within microcolonies

When analyzing the location of reporting cells in microcolonies, we noticed that cells in the center of the microcolonies, corresponding to the location with the highest cell stacking, were among the first to activate *PmnxG* (**Fig. 3a**), followed within 1 h by detectable fluorescence in the outer rim of the microcolony (**Fig. 3a, c**). About 2.5 hours later, the bioreporter signal appeared everywhere in the microcolony. The average rate of increase in the *PmnxG* bioreporter signal was fastest at the microcolony edges (**Fig. 3c**). These observations suggest that cells in the outermost layer of the microcolony experience optimal conditions for promoter activation. Reporter activation in the *PmcoA* strain started 5 h later than in the *PmnxG* strain (**Fig. 3b**), with a pattern that radiated from the center outwards but never reached the edge of the microcolonies (**Fig. 3b** and **d**). These reporter expression patterns were observed consistently among different microcolonies, suggesting that the activation of the Mn oxidase gene promoters results from gradients in chemical cues across the microcolonies.

**Fig. 3.**
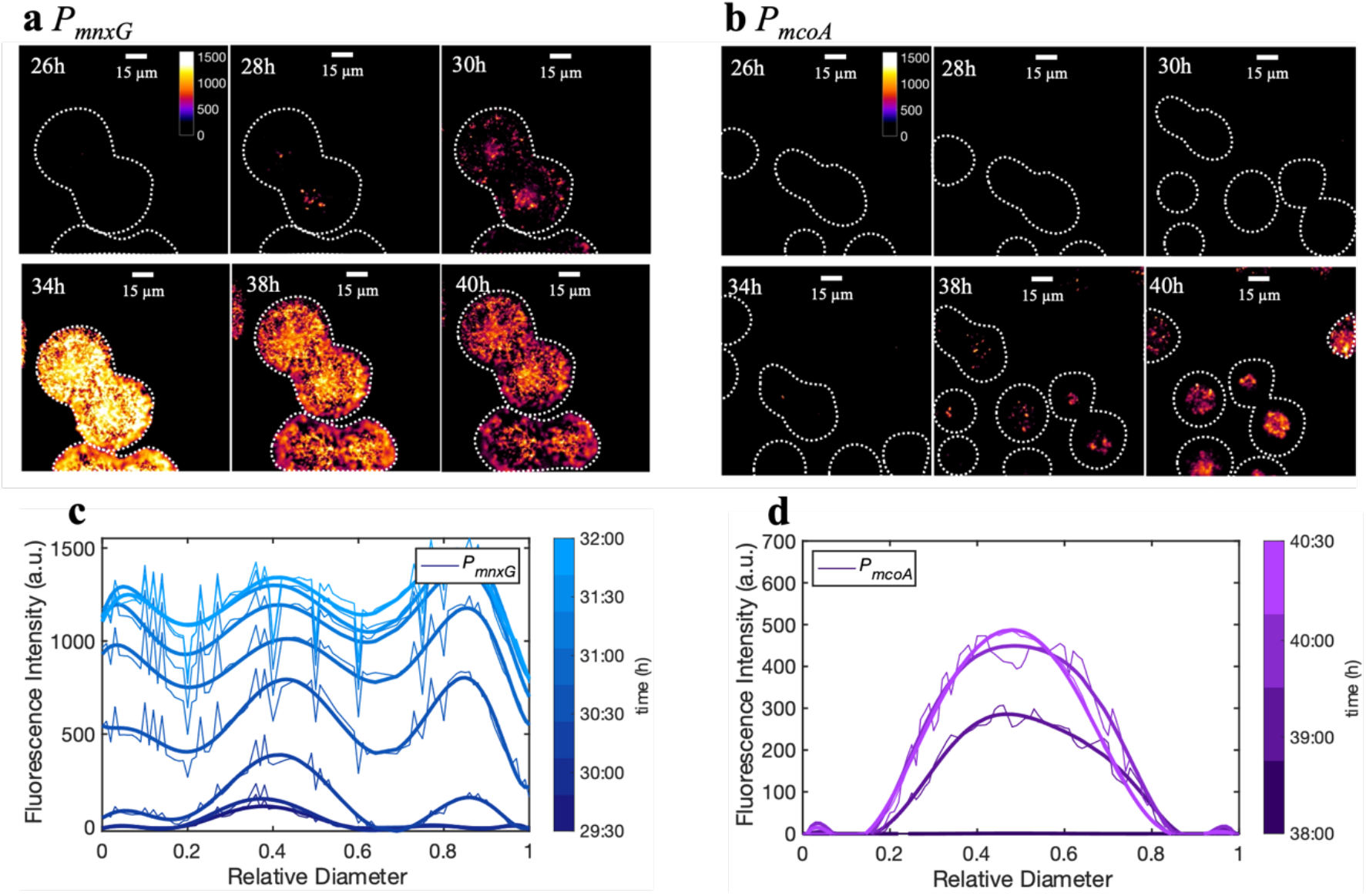
Spatiotemporal activation of Mn oxidase gene promoters of *P. putida* GB-1 in surface-grown microcolonies. Microcolonies of *PmnxG* (**a**) and *PmcoA* (**b**) bioreporter strains imaged at different times after entry into stationary phase. The fluorescence signal is shown as a heatmap, with values defined as the signal above the threshold. The threshold is the 99th quantile of wild-type GB-1 fluorescence distribution in the reporter wavelengths, calculated using background-subtracted images. The dotted lines show the boundaries of the microcolonies obtained from phase contrast images. Note the later activation of *mcoA* compared to *mnxG*. The decrease in fluorescence intensity for *PmnxG* after 34 h is due to photobleaching (**SI Appendix, Fig. S7**). **c-d** Fluorescence intensity profiles normalized by the microcolony diameter during expression of *PmnxG* (**c**) and *PmcoA* (**d**). Profiles represent the average of 6 replicates for each strain; trends are shown by smoothing using a polynomial. Note the edge appearance of *mnxG* promoter fused fluorescent protein and center appearance of *mcoA*.

To verify that the spatiotemporal patterns we observed did not result from the restrictive growth conditions caused by the microscope chambers (i.e., low oxygen availability, **Supplementary** Fig. 10), we monitored *PmnxG* and *PmcoA* activation in open-chamber experiments, where the coverslip was removed to allow for maximum airflow (**Supplementary** Figs. 10 and 11). We found the same patterns of promoter activation in microcolonies grown in both closed and open chambers (**Supplementary** Fig. 11). Notably, the proportion of cells activating *PmcoA* increased with increasing microcolony size (**Supplementary** Fig. 12), and *PmcoA* was expressed earlier in the center of cell flocs than in single planktonic cells in liquid culture (**Fig. 2d**), suggesting more favorable conditions or cues that could activate the *PmcoA* promoter in dense multicellular aggregates.

### Activation of the Mn oxidase gene promoters correlates to the formation of Mn oxide precipitates

To confirm that the bioreporter signal is a faithful representation of the onset of Mn biomineralization, we compared the timing and the extent of reporter fluorescence with the formation of Mn oxide precipitates. Indeed, visible Mn oxide precipitates appeared within an hour of *PmnxG* activation in both stationary phase microcolonies (**Fig. 4a**) and liquid-suspended culture (**Fig. 4d**). While liquid-suspended cultures showed a small amount of Mn removal from solution prior to promoter activation (2.3 ± 0.9 µM Mn(II), **Fig. 4d**), this loss of Mn from solution resulted from sorption of Mn(II) by the biomass rather than its enzymatic oxidation to Mn(III, IV) (**Fig. 4d** and **Supplementary** Fig. 13).

**Fig. 4.**
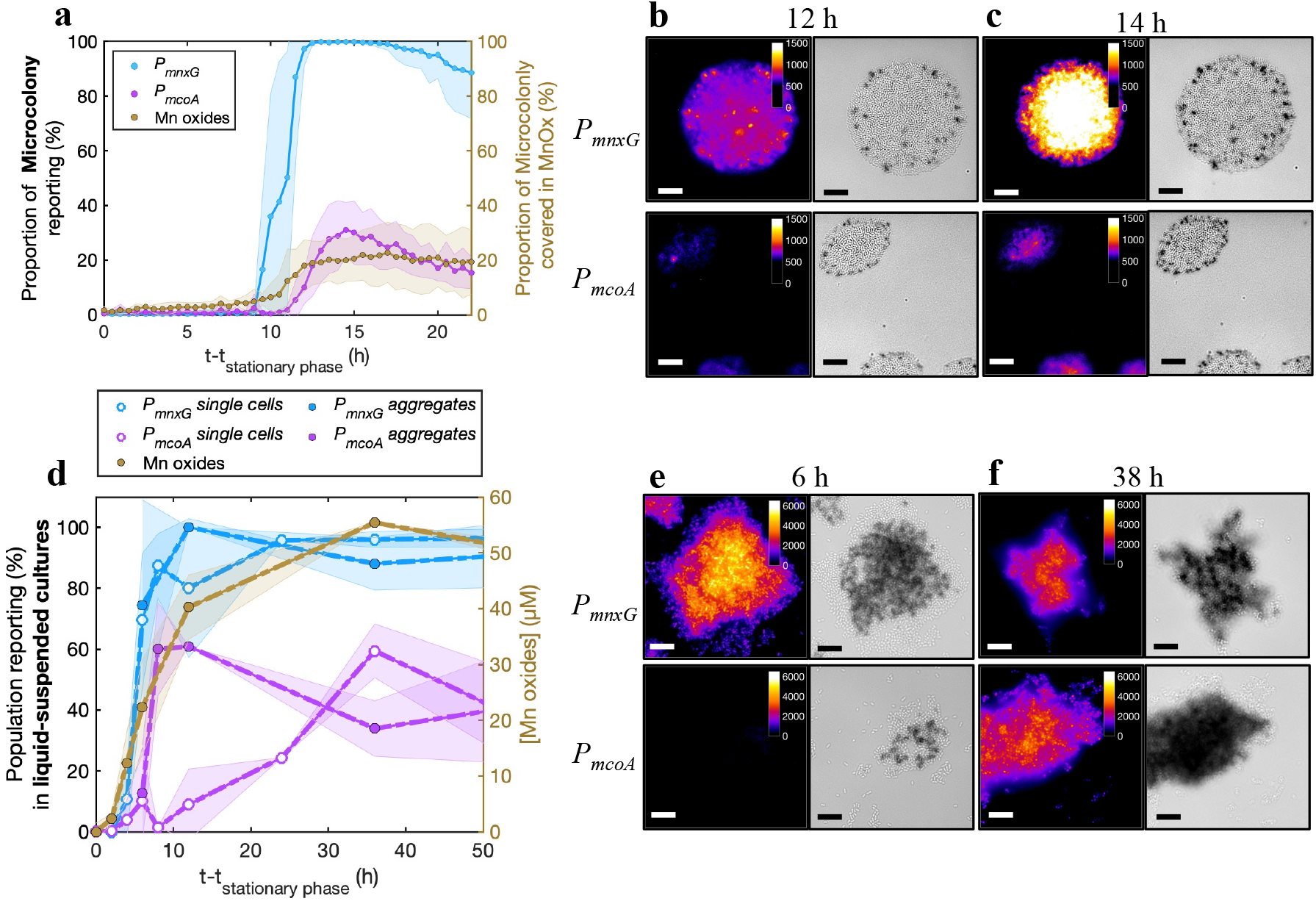
Correlation between *mnxG* and *mcoA* promoter activation and Mn oxide formation in *P. putida* GB-1. **a** Proportion of microcolony area with activated bioreporter fluorescence signal (left axis, above threshold; *mnxG* in blue, and *mcoA* in magenta), and the proportion of microcolony area covered with visible Mn oxide precipitates (right axis, in brown). The time axis is presented as time in (h) after the onset of the stationary phase (reached after 27.5 h on a solid surface). Lines connect the means, and transparent areas represent ± one standard deviation (n = 20 colonies analyzed for each reporter). **b-c** Representative microscopy images of *PmnxG* and *PmcoA* bioreporter microcolonies 12 h **(b**) or 14 h (**c**) in stationary phase for intensity of the eCherry fluorescence (left, according to color scale) or bright field (corresponding Mn oxide precipitates; in gray). Scale bars denote 10 µm. **d** Proportion of activated fluorescent reporter (left axis); *mnxG* in blue and *mcoA* in magenta) as either individual planktonic cells (open circles) or in aggregates (filled circles) in liquid suspended culture, and corresponding fraction of Mn oxides (in µM, right axis) over time after entry into stationary phase (12 h after culture start). The amount of Mn oxide precipitated was calculated by subtraction of the aqueous Mn to the total Mn, as measured by ICP-MS. Dotted lines connect sample means with transparent colored areas representing ± one standard deviation (n = 3 replicates). **e-f** as **b** and **c,** but for aggregates from liquid suspended culture after 6 h (**e**) and 38 h (**f**) into stationary phase. Scale bars represent 10 µm.

Microcolonies grown under closed chamber conditions typically showed a strong correlation between hotspots of promoter activation and Mn-oxide precipitates (**Fig. 4b** and **Supplementary** Fig. 14). These Mn oxide hotspots were concentrated around the edge of the microcolonies (**Fig. 4b**, **c** and **Supplementary** Fig. 14), which coincides with the location of early *PmnxG* activation (**Fig. 3a**). Despite the increase in the proportion of *PmnxG* activation to up to 99.6% ± 0.7% of the microcolonies, Mn oxide precipitates covered only 22.7% ± 2.8% of the microcolony surface after 24 h in stationary phase (**Fig. 4a**). Since the bioreporter strains have wild-type GB-1 background, microcolonies of the *PmcoA* bioreporter also showed Mn-oxide precipitates near the microcolony edges (**Fig. 4b, c**), but these precipitates did not coincide with *PmcoA* activation. Reporter fluorescence from *PmcoA* appeared later and in the center of the microcolonies (**Fig. 4a-c**) but did not correlate to a further increase in Mn oxide precipitation (**Fig. 4a-c**). Together, these results demonstrate that MnxG was responsible for the initial formation of Mn-oxides. However, the sparse surface coverage with Mn oxide in other areas of the microcolonies under continued expression of *mnxG* and later expression of *mcoA* suggests that other requirements for mineral precipitation were not fulfilled in the closed chambers.

When experiments were repeated under open chamber conditions in wild-type GB-1, the projected microcolony surface area was fully covered with Mn oxide precipitates (**Supplementary** Fig. 15a), coinciding with the reporter gene expression observed in the closed chambers (**Supplementary** Fig. 11). Similar culturing of a GB-1 strain deleted for *mcoA* showed most Mn oxides near the edges and less in the colony center, whereas the GB-1 strain deleted for *mnxG* only formed Mn oxides in the colony center (**Supplementary** Fig. 15a, b). These patterns of Mn oxide formation are similar to the global localization of reporter fluorescence from each Mn-oxidase-specific gene promoter. The presence and sequential activation of two Mn oxidases in GB-1 thus expands the environmental conditions and the time window for Mn biomineralization. Additionally, the reason for the sparser coverage of the microcolonies with Mn oxides in the closed chamber configuration, despite expression of the Mn-oxidase gene promoters, is likely due to the lower oxygen flux towards the cells^49^, which would deprive the enzymes of one of their co-substrates. This oxygen limitation in the closed chambers was removed the open chamber experiments where the microcolony size increased 10-fold and Mn oxides covered the entire microcolony surface.

In liquid-suspended culture, we also observed a strong correlation between fluorescent signal appearance and the onset of Mn oxide precipitation for the *PmnxG* reporter strain (**Fig. 4d, e**). The formation of Mn oxides around cells and cell aggregates that activated the *PmnxG* promoter, but had not yet expressed *mcoA* (PmcoA activation appeared ca. 6 h later, **Fig. 4e**), confirms that MnxG is sufficient for initial Mn oxidation and precipitation (**Fig. 4e**). By chemically measuring the extent of Mn oxide precipitation, we found that wild-type GB-1 and the *mcoA* deletion strain remove aqueous Mn(II) from solution at the same rate (ca. 0.41 h^−1^), whereas the *mnxG* deletion strain shows an order of magnitude slower rate (0.04 h^-1^, **Supplementary** Fig. 16). Finally, no Mn oxide precipitation was observed in the double knockout strain under the tested conditions (Supplementary Fig. 16). This confirms that MnxG is the main Mn oxidase, followed by McoA, whereas MopA activity is not present.

## Discussion

Microbial Mn oxidation was discovered a century ago^50^. This process is performed by a large number of phylogenetically diverse bacteria and fungi^5,15^, yet the environmental controls on biomineralization and its physiological function remain elusive^17^. Using fluorescent gene reporters to target the activation of the promoters upstream of *mnxG* and *mcoA* in *P. putida* GB-1, we provide the first temporally and spatially resolved analysis of Mn oxidase promoter activation. We show that reporter activation coincides spatially with Mn oxide precipitation in microcolonies and cellular aggregates, demonstrating that reporter signal can be used to study Mn oxidation at the individual cell level. This approach has provided new insights regarding the regulation of bacterial Mn oxidation, which notably occurs only in non-dividing stationary phase cells and only in the presence of Mn, is confined to subsets of cells within the population, and is different for both Mn oxidase genes.

Previously, Mn biomineralization has been studied based on the extracellular appearance of Mn oxides ^5,26,29,33^ or by enzyme purification^22^, leading to the hypothesis that the presence of multiple Mn oxidases is linked to differences in lifestyle (e.g., biofilm or planktonic cells)^26,31^. Here, we find no evidence for exclusive expression of either *mnxG* or *mcoA* in sessile (microcolonies) or planktonic cells. Instead, *mnxG* and *mcoA* are expressed under both growth conditions. Manganese oxidase gene activation has never been attributed to a specific growth phase, but we can now show beyond doubt, from both surface-grown and liquid-suspended culture experiments, that *mnxG* and *mcoA* are activated during stationary phase conditions and only in the presence of aqueous Mn (**Fig. 1** and **Supplementary** Fig. 5). The requirement of Mn for promoter activation corroborates the finding that additional specific regulatory factors, such as the proposed MnxS1/S2 sensor histidine kinases and MnxR protein^27^ are needed to initiate *mnxG* and/or *mcoA* expression^51^.

Despite the expression of both *mnxG* and *mcoA* during stationary phase, their activation was non-uniform in both surface and liquid-grown cells. First, the proportion of the population expressing either of these genes (by the proxy of the promoter fusion to the fluorescent protein) increased over time for both promoters, with a higher proportion of the population activating *mnxG* than *mcoA*. This bimodal gene activation confers significant phenotypic heterogeneity to individuals within the population of *P. putida* GB-1 cells. Second, the onset of promoter activation relative to entry into the stationary phase differed between the two genes, with *mnxG* being expressed earlier than *mcoA*. Third, the localization of cells expressing either *mnxG* or *mcoA* differed within stationary phase colonies, with *mnxG* expression starting from the edges and moving inward over time, and *mcoA* more confined to colony centers. These observations show that *mnxG* and *mcoA* respond to different chemical gradients forming across the colonies and behave synergistically.

Population-level controls over bacterial gene activation have been attributed to mutation, stress response, intra-population dynamics (e.g., quorum sensing), or variation in chemical conditions (e.g., microenvironments), amongst other factors^47,48,52–54^. The high reproducibility between biological replicates, the onset of expression in non-growing cells, and the observed loss of bioreporter signal in exponentially growing cells suggest that the observed bimodality is not the result of a reproducible genetic switch (e.g., phase variation), but is rather a response to environmental cues. Sequential activation of *mnxG* and *mcoA* might result from changes in extracellular conditions, such as pH^55,56^ or Eh^57^, or secondary metabolites^58,59^, which often evolve during bacterial growth in solid and liquid media^60,61^. The delayed activation of *mcoA* relative to *mnxG* and the difference in the timing of *mcoA* activation in cells grown on solid surfaces and liquid-suspended cultures suggests that *mcoA* is more sensitive to environmental conditions encountered later in stationary phase and prevalent in the colony or aggregate center (**Fig. 5**). The most common dynamic gradient within microcolonies involves oxygen concentration^49,62–65^. During active colony expansion, oxygen consumption near the edge of the colonies leads to its depletion inside the colony, as observed by Díaz-Pascual *et al*.^64^. However, once the cells reach stationary phase, they consume less oxygen and oxygen is replenished towards the colony center within ca. 8 h^64^. The timing of oxygen depletion and renewed diffusion within microcolonies observed by Díaz-Pascual *et al*.^64^ are remarkably similar to the *mnxG* and *mcoA* promoter activation patterns in GB-1 reporter cells. This suggests that oxygen is a co-substrate for Mn oxidase gene activation in stationary phase, and its flux or local concentration determines expression onset. In this scenario, while the oxygen inflow is replenished to the microcolony edges, *mnxG* is activated, and as oxygen diffusion proceeds to the colony center, *mcoA* becomes activated (**Fig. 3** and **Supplementary 11**). The specific consumption of oxygen associated with MnxG activity and precipitation of Mn oxides may further delay the onset of *mcoA* expression or alternatively trigger *mcoA* activation, if the latter operates at a lower oxygen threshold. The earlier activation of *mcoA* in the center of the cell aggregates compared to single cells (**Fig. 2d**) confirms that the local conditions created in the center of the aggregates, which are commonly oxygen-limited^66,67^, favor activation of *mcoA* (**Fig. 5**). Future work can now explore how relevant chemical gradients (e.g., Mn, O2, secondary metabolites, and signaling molecules) and their interactions regulate the heterogeneous expression of the Mn oxidases and subsequent mineral precipitation.

**Fig. 5.**
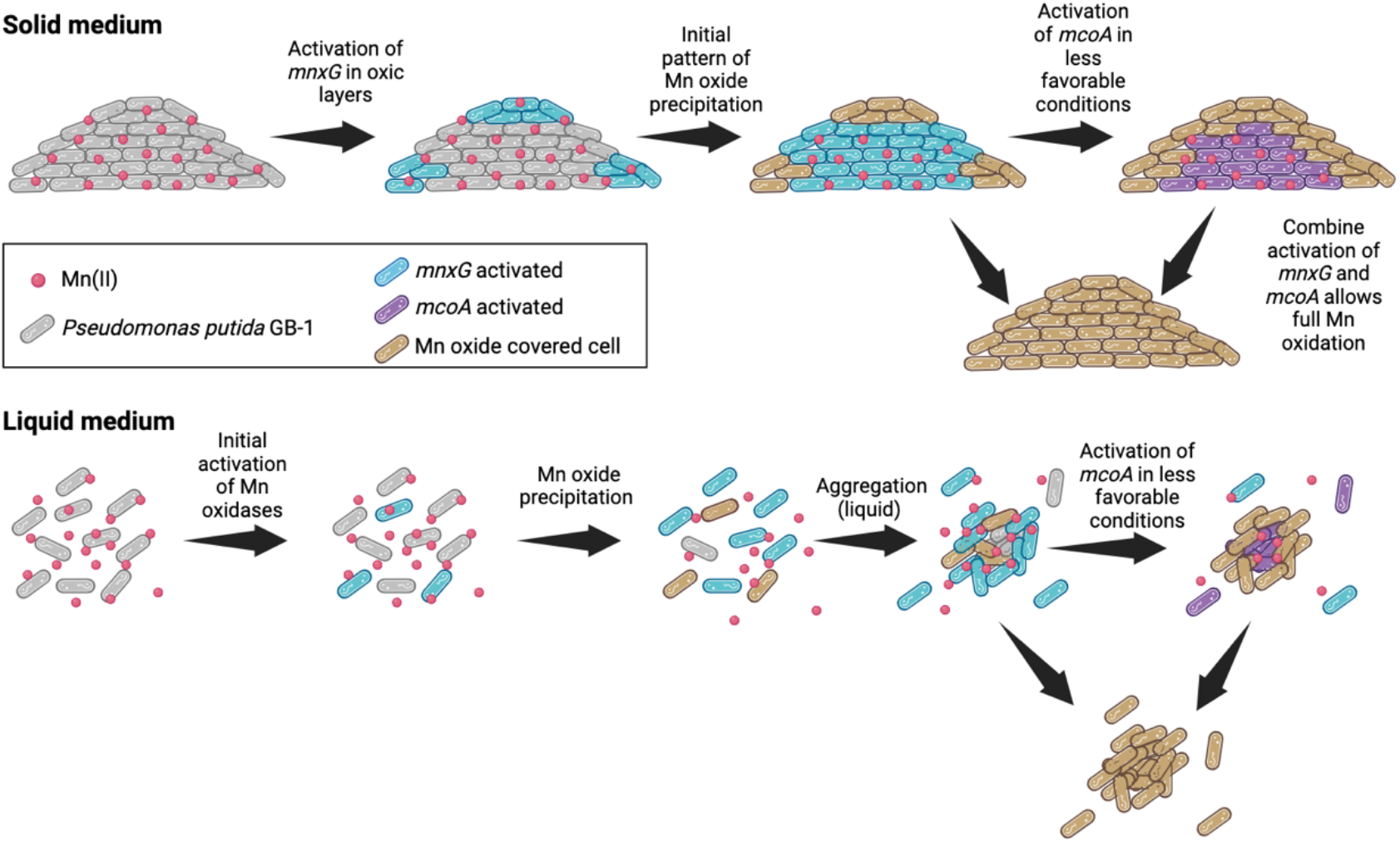
Proposed mechanism of promoter activation and Mn oxide biomineralization in *Pseudomonas putida* GB-1. The cells in microcolonies grown to stationary phase in the presence of Mn(II) activate the primary pathway, *mnxG*, under favorable conditions for Mn oxidation (i.e. high O2 and circumneutral pH). The microenvironment that develops at the center of the microcolony creates less favorable conditions for Mn oxidation, leading to the activation of the second Mn oxidase, *mcoA*. The combined activation of the two enzymes leads to full Mn oxidation and its removal from the solution to the solid phase as Mn oxide. In liquid, the same mechanism is observed, where the aggregate core has less favorable conditions for Mn oxidation, leading to the activation of *mcoA* in the center. The activation of *mcoA* in single cells likely results from delayed changes in solution chemistry.

The preferential activation of *mnxG*, as deduced by the earlier timing, broader spatial extent, and occurrence in a higher proportion of the population, concomitant with the precipitation of Mn oxide in all the conditions tested (i.e., closed chambers, open chambers, and liquid medium), demonstrates that MnxG is the dominant oxidase for *P. putida* GB-1. This is supported by experiments with GB-1 strains lacking either *mnxG* or *mcoA*, which show that MnxG alone can oxidize the totality of the Mn(II) supplied (**Supplementary** Fig. 16), and that McoA is less efficient and remains largely complementary (**Figs. 4** and **5**). Overall, our results indicate that activation of *mcoA* following activation of *mnxG* can drive Mn oxidation under sub-optimal conditions for MnxG or when its functional capacity is exceeded (**Fig. 5**).

By carrying multiple Mn oxidases, as regularly found in Mn oxidizers^15^, GB-1 can perform Mn oxidation under a wider range of environmental conditions. Bimodal activation of *mnxG* and *mcoA* would lower the cost of producing energetically expensive enzymes in all cells while sharing the benefit of Mn oxide precipitation and/or aqueous Mn(II) removal with the whole population or community. This could be considered a microbial cooperation strategy, as proposed for other metabolic functions, that provides a competitive or ecological advantage^52,68–70^. Several hypotheses for the advantage of Mn oxidation for bacteria have been put forward, but none have been firmly demonstrated. For example, Mn oxidation may provide a pathway for metal homeostasis through increased expression of proteins that allow for sequestration of Mn in the solid phase^71^, enable the generation of bioavailable carbon substrates from the reaction of Mn oxide and complex organic matter^38^, or allow control over extracellular redox conditions to support cell maintenance at stationary phase^72^. Whether this stationary phase process is initiated to rid the extracellular environment of aqueous Mn or to promote the formation of the manganese oxides requires further study. Our discovery that social cooperation within a bacterial population may underlay the formation of biominerals provides a new framework to investigate the function of Mn oxidation and biomineralization not only for individual cells, but for microbial communities and ecosystems.

## Materials and Methods

### Bacterial strain and plasmid construction

The strains used in our study are derivatives of *P. putida* GB-1 (**Supplementary Table 1**). *Escherichia coli* DH5alpha was used to propagate plasmid DNA, which was used as a vector for the construction of the reporter gene fusions. All enzymes used for DNA digestion or ligation were purchased from New England Biolabs. PCR assays to amplify DNA were performed with primers described in **Supplementary Table 2** following the manufacturer’s instructions. DNA and PCR products were purified using Nucleospin Gel and PCR Clean-up kits (Macherey-Nagel) according to the manufacturer’s instructions.

The promoter regions upstream of *mnxG* (*2447*) and *mcoA* (*2665*) were amplified with genomic DNA of *P. putida* GB-1 as a template, using reverse (R) and forward (F) primer pairs PputGB1_2447.F/PputGB1_2447.R and PputGB1_2665L.F/ PputGB1_2665.R, respectively (**Supplementary** Figs. 1 and 2 and **Table 2**). The reporter gene *echerry* (GenBank accession number: AY678264) and its ribosomal binding site (RBS) were amplified on the plasmid DNA pMQ64-echerry^73^ using the primer pair eCherry.F/ eCherryTn5.R (**Supplementary Table 2**). Each promoter region was placed upstream of the *echerry* gene and cloned into SmaI-digested pBAM1^74^, using the ClonExpress II one step cloning kit (Vazyme). This produced plasmids pBAM1(miniTn5::*PmnxG-echerry*), and pBAM1(miniTn5::*PmcoA-echerry*). The resulting plasmids were verified by restriction profiling and DNA sequencing. Purified plasmid DNA was then introduced into *P. putida* GB-1 by electro-transformation, and clones with a single integrated copy of the mini-transposon reporter construct were selected^74–77^. Three independent clones of *P. putida* GB1 with potentially different integration sites of the reporter constructs were purified and stored at –80 °C. Growth rates, fluorescence intensity, and Mn oxide precipitation (rate, onset, and time to full oxidation) were tested for each clone and compared to the wild type (**Supplementary** Fig. 17 and **Table 4**). Electroporation was carried out as described by Dower *et al*.^78^, in a Bio-Rad GenePulser Xcell apparatus set at 25 µF, 200 V and 2.5 kV for *E. coli* and 2.2 kV for *P. putida,* using 2-mm gap electroporation cuvettes (Cellprojects). Plasmid or insert DNA sequencing was performed by MycroSynth (Switzerland).

### Growth medium and inoculum

*P. putida* GB-1 bioreporter strains were grown for 16 h in Luria Broth (LB) containing 25 µg ml^-1^ of kanamycin at 30°C in an orbital shaker at 180 rpm. Cells were then centrifuged at 4000 x g for 1 min and resuspended in MSTA salt solution (**Supplementary Table 3**), at room temperature. This step was repeated three times to wash the cell suspension and remove the spent growth medium. To diminish aggregation of *P. putida* GB-1 cells in Mn oxidation studies, as is commonly observed on complex media (e.g. glucose, casamino acids, yeast extract)^26,29,33^, we developed a defined growth medium for Mn oxidation, hereafter MSTA (**Supplementary** Fig. 18 and **Table 3**). Briefly, the medium employed in this study contained 0.4 mM CaCl2·2H2O, 0.25 mM MgSO4·H2O, 0.25 mM Na2HPO4, 0.15 mM KH2PO4, 1:2 Fe(III)-EDTA (prepared by reacting 20 µM FeCl3·6H2O with 40 µM EDTA, adjusted at pH 6.5 with NaOH), HEPES buffer (prepared by adjusting the pH to 7.0 with NaOH), 5 mM (NH4)2SO4, 40 nM CuSO4·5H2O, 273 nM ZnSO4·7H2O, 84 nM CoCl2·6H2O, and 53.7nM NaMoO4·2H2O and 0.5 mM L-arginine as the carbon source. The growth medium was complemented with either 0 or 50 µM Mn(II) added as manganese chloride (MnCl2). All cell cultures were grown in sterile Erlenmeyer flasks with a 2:5 liquid-to-air ratio, to maintain sufficient oxygenation, at 30°C and orbital shaking at 180 rpm, in the dark. The washed cell suspension was transferred to MSTA medium at a starting optical density (OD600) of 0.01.

### Gene expression and Mn oxidation in microcolonies

Microscopy chambers (Helmut Saur Laborbedarf, Germany) were used to follow the growth of single cells over time, the appearance of the fluorescence bioreporter signal, and Mn oxide coverage (**Supplementary** Fig. 19). Cells were seeded on agarose patches^79^, made using the MSTA medium and supplemented with 1% agarose, unless specified otherwise. Then, 3 µl of the washed and diluted cell suspension (OD600nm of 0.01) was deposited onto the agarose patches and sealed into the microscopy chamber. To allow for airflow and oxygenation, we punctured the silicone gaskets on opposite sides, leaving two needles in place throughout the experiment. The microscopy chamber was then mounted on the microscope, equipped with a temperature-controlled incubator maintained at 30°C. Epifluorescence microscopy images were acquired every 30 min for 48 h or 54 h. Additional experiments where the microscopy chambers were kept uncovered during bacterial growth and Mn oxidation were conducted in the same way and are referred to as open chamber experiments.

### Gene expression and Mn oxidation in liquid-suspended cultures

Triplicate cultures for each of *P. putida* GB-1 wild-type strain, *PmnxG* strain, and *PmcoA* strain were grown at 30°C and 180 rpm. Up to eight samples were collected separately for epifluorescence microscopy imaging and ICP-MS measurements between 12 and 72 h. For imaging, three microdroplets (4 µl) were deposited on a coated glass slide. To prepare coated slides, 600 µl of a 1% agarose and MSTA salt solution was deposited on the glass slide, which was covered with a second glass slide and allowed to cool before removing the top slide. For Mn quantification in solution and in the solid phase, ICP-MS analysis was performed on filtered or acid-digested aliquots, respectively (see **Supplementary Material and Methods**). Our method has a limit of quantification of 0.04 µg L^-1^ or 0.67 nM.

### Epifluorescence microscopy

*P. putida* cells were imaged at 1000x magnification, using a Nikon Eclipse Ti2 equipped with a Hamamatsu ORCA-Flash 4.0 camera, a Lumencor Light Engine LIDA 3-color light source, and SOLA III solid-state white light excitation source. Images were acquired in 2048 by 2044 field of view and a pixel resolution of 0.07 µm/pixel. Samples from liquid-suspended cultures were imaged at the edge of the sample to capture planktonic cells and randomly near the center to capture both planktonic and aggregated cells. Cells were imaged in phase contrast (10 ms exposure). eCherry fluorescence was captured by excitation at 562 nm using a SOLA III light engine at 50% intensity and recording emission at 645.5 ± 50 nm (500 ms exposure). Images were stored as 16-bit TIFF files.

### Image analysis

Cell aggregates and the microcolonies were analyzed using MATLAB R2021b scripts, developed and adapted to extract the boundaries of the microcolony or aggregate and the corresponding fluorescence signal (**Supplementary Material and Methods** and **Supplementary** Fig. 20). In the case of single (non-aggregated) cells on images, we used SuperSegger for segmentation^80^ (**Supplementary Material and Methods** and **Supplementary** Fig. 21). The mean cell fluorescence values (sum of the fluorescent pixels normalized by the cell area) or pixel fluorescence (for aggregates and microcolonies) were corrected by subtracting the median background signal outside the segmented cells. Images in the eCherry channel of wild-type GB-1 cells were used to identify the cell auto-fluorescence, which was defined as the 99^th^ quantile of the wild type fluorescence distribution and used as the threshold above which we considered a ‘true’ eCherry bioreporter signal. Similarly, the quantification of Mn oxide precipitates on microcolonies was performed using MATLAB scripts developed to identify the proportion and intensity of the color change given by the appearance of Mn oxides (**Supplementary Material and Methods** and **Supplementary** Fig. 22).

## Statistics

All statistical analyses were performed using MATLAB (v. 2024a). Outliers were removed at 6σ for visualization purposes. The proportion of the population reporting included all of the data, calculated as the number of cells with fluorescence values above the 99^th^ quantile of the wild type fluorescence distribution. All experiments were performed in three to five replicates. Each separate microcolonies were considered as replicates and represented 7 to 21 replicates for the conditions in the absence of Mn across all experiments. In its presence, the number of replicates represented 11 to 45 replicates.

## Data and code availability

All data presented in this work are available within the article and the supplementary files. Source data and codes can be found on Dryad data repository at DOI: 10.5061/dryad.fj6q5742z. Any additional requests can be addressed to the corresponding authors.

## Acknowledgments

We thank Dr. Vladimir Sentchilo for his help in the development of the bioreporters, Dr. Kyounglim Kang for her assistance with ICP-MS, Tania Miguel Trabajo for providing training in the use of the sealed microscope chambers, and Konane Gurfield for her help with image collection. We are also grateful to Eleanor Fadely and Dr. Kyounglim Kang for their helpful discussions. This work was supported by the Swiss National Science Foundation (200021_188546) and the U.S. National Science Foundation (1449501 and 2322428). The funder played no role in study design, data collection, analysis and interpretation of data, or the writing of this manuscript.

## Author Contributions

G.G designed the research, performed the research, analyzed the data, and wrote the paper; N.C. designed the research, performed the research, analyzed the data; J.R.V.D.M. designed the research and wrote the paper; J.P. designed the research, analyzed the data, and wrote the paper.

## Competing Interest Statement

All authors declare no financial or non-financial competing interests.

## Notes

### Competing Interest Statement

The authors have declared no competing interest.

