## Supplemental Figures 1 to 22 and tables 1 to 4 for "Population-level control of two manganese oxidases expands the niche for bacterial manganese biomineralization"

Jasquelin Peña

This PDF file includes:

Supplementary information (SI) text

Supplementary Figures 1 to 22

Supplementary Tables S1 to S4

SI References

### Supplementary Information Text

#### Supplementary Material and Methods

**Plasmid and strain storage information.** *Escherichia coli* DH5αlpir with kanamycin antibiotic resistance, obtained from the bacterial library of Jan van der Meer, was used as the host strain for the plasmid construction of the $P_{mnxG}$ and $P_{mcoA}$ bioreporters. *E. coli* was grown in LB medium supplemented with 20 µg/ml kanamycin at 37°C and shaking at 180 rpm. The strains, stored at -80°C in 20% glycerol and 80% LB, were grown on LB plates with 25 µg/ml of kanamycin. An isolated colony was picked and inoculated in liquid LB containing the same antibiotic concentration and grown until the stationary phase.

**Construction of the bioreporter strains.** The promoter region for $P_{mnxG}$ was selected as the 266 base pairs in front of the gene *mnxG* (MnxG) and the promoter region for $P_{mcoA}$ was selected as the 534 base pairs in front of the gene *mcoA* (McoA), identified by Geszvain et al.[1] in *Pseudomonas putida* GB-1. The promoter sequences and eCherry sequence (711 base pairs), contained in pMQ64eche from *E. coli* DH5α with kanamycin resistance, were amplified by PCR (with denaturation at 98°C for 30 sec, 30 cycles of amplification with 10 sec at 98°C, 30 sec at 67°C, and 72°C for 3 min) with the corresponding primers (see **Supplementary Table 2**). We checked the purity of the PCR products by gel migration and purified the PCR products using a NucleoSpin® Gel and PCR clean-up Kit (Ref: 740609.50) from Macherey-Nagel. To construct the $P_{mnxG}$_eCherry and $P_{mcoA}$_eCherry plasmids, we extracted the pBAM1 plasmid, contained in *E .coli* DH5α, using a NucleoSpin® Plasmid extraction kit from Macherey-Nagel (Ref: 740588.250) for low-copy plasmid extraction. We then linearized pBAM1 with a SmaI digestion and, to prevent unwanted recombination, digested the BamHI site to break down the recombinant DNA fragment. The linearized pBAMI was then mixed with *P. putida* genomic DNA ($P_{mnxG}$ or $P_{mcoA}$) and the eCherry vector for 30 min at 37°C. The recombined circular plasmid DNA was then transformed into chemically competent *E. coli* DH5α. We selected four biological replicates from the viable transformed colonies that grew on LB Petri dishes containing kanamycin 25 $\mu g \cdot ml^{-1}$ and confirmed the construction of the plasmid by sequencing.

**Plasmid transformation into *Pseudomonas putida* GB-1.** We prepared electrocompetent *P. putida* GB-1 cells by washing three times a 1.5 ml aliquot of cell suspension with MOPS-Glycerol. The suspension was centrifuged at 6000 x g for 1 min at 4°C. The supernatant was then discarded and the pellet was resuspended in MOPS-Glycerol. After the third washing, the pellet was resuspended in 50 µl MOPS-Glycerol. We added 3 µl of previously extracted plasmids containing $P_{mnxG}$ or $P_{mcoA}$, at a concentration of 40-60 $ng \cdot \mu l^{-1}$, and transferred 53 µl of the suspension into 2 mm electroporation cuvettes. We then injected a pulse of current of 2.2 kV (capacitance 25 mF, resistivity 200 Ω) using a BioRAD Gene PulserXcell™. The resulting cell suspensions were then resuspended in 600 µl S.O.C. medium and incubated at 35°C for 1h40 and at room temperature for another 2h. To select the transformed stains, we transferred the cultures onto LB Petri dishes containing 25 $\mu g \cdot ml^{-1}$ of kanamycin. Five clones were selected from the viable colonies and stored in liquid LB with 20% glycerol at -80°C.

**Single and double knockout mutants.** *mnxG* and *mcoA* single deletion mutants, and a double knockout mutant, were constructed using the two-step chromosomal gene inactivation technique, as described elsewhere[2,3] and using the primers described in **Supplementary Table 2**. We confirmed the deletion of *mnxG* and *mcoA* by PCR amplification (**Supplementary Table 2**). The defect of the double knockout mutant on Mn oxidation was confirmed by ICP-MS analysis (**Supplementary Fig. 16**).

**Testing the growth, fluorescence, and Mn oxide yield of the transformed clones by spectrofluorometry.** Optical density (OD) and fluorescence intensity were measured with a Varioskan LUX plate reader (Multimode Microplate Reader) using transparent plastic 96 well plates of 350 µL well capacity. A volume of 200 µL of sample was used to promote oxygenation and mixing. The temperature was maintained at 30°C and agitation was performed under continuous double orbital shaking at 425 rpm. The OD was measured at 600 nm. The fluorescence was measured in the mCherry channel with excitation at 579 nm and emission at 616 nm. A single measurement of OD and fluorescence consisted of an average of 8 measurements in the center of the well with a frequency of 100 ms at 7 mm above the well using a Xenon Flashlight source at high energy. Each measurement was conducted in six replicates. Growth rates were calculated assuming first-order kinetics during the exponential growth phase. The clones were tested against the wild-type to confirm that the Tn5 random insertion did not disturb bacterial growth and/or ability to precipitate Mn oxide. The five clones were also tested against each other to confirm reporter gene activation. To further confirm that the Tn5 insertion did not affect the Mn oxidation capacity, we systematically compared the Mn(II) oxidation kinetics of the bioreporters against the wild-type and did not find any significant difference (**Supplementary Table 4**).

**Viability of reporting cells on agarose surfaces.** To test for the viability of the subpopulations showing $P_{mnxG}$ or $P_{mcoA}$ activation, we first grew the cells in liquid MSTA medium (**Supplementary Table 3**) containing 5 mM L-Arginine and 50 µM $MnCl_2$. After 20h (when $P_{mnxG}$ is active, but $P_{mcoA}$ is not), we washed the cell suspension in MSTA salts and inoculated 1% agarose patches containing 5 mM L-Arginine and MSTA salts, in the absence of $MnCl_2$. Reporting and non-reporting cell growth was then tracked using Dimalis, an image segmentation tool developed for microcolony growth segmentation[4].

**Development of the MSTA growth medium.** To optimize Mn oxide precipitation while lowering the cell suspension aggregation, we developed a defined growth medium based on Leptothrix and MSTP media. *Pseudomonas putida* GB-1 were grown in Lept medium salts with combinations of casamino acids, yeast extract, and glucose (**Supplementary Fig. 18**). The Mn oxides within the liquid-suspended cultures were quantified by ICP-MS after acid digestion. The optimal carbon source was determined as casamino acids. This carbon source was then refined to select single amino acids leading to Mn oxide precipitation. Next, 5 mM of each amino acid was added to MST salts[5,6] and the Mn oxide yield was assessed with LBB (**Supplementary Fig. 18**). L-arginine was selected as the best candidate due to the low aggregation, precipitation of Mn oxides, and high growth. The composition of this growth medium, MSTA, is described in **Supplementary Table 3**.

**Mn(II) sorption into the biomass.** Sorption experiments were conducted in MSTA containing 50 µM $MnCl_2$, in triplicates. The bioreporter strains were grown for 48h and aliquot samples were taken during late exponential and stationary phases at 10, 12, 14, 16, 18, and 21 h. Total Mn was measured by ICP-MS on samples obtained by digesting 1 ml of the biomass suspension in 0.5 ml nitric acid 65% and 0.5 ml oxalic acid 0.4 M. The aqueous Mn was measured by ICP-MS on the

0.22 μm filtered samples. The sorbed Mn is reported as the difference between the total Mn and the aqueous Mn (**Supplementary Fig. 15**). Finally, to confirm that the discrepancy between the total and aqueous Mn originated from the sorbed fraction, we monitored the presence of Mn oxides using the LBB redox dye[7] (**Supplementary Fig. 15)**.

**Image analysis of microcolonies and aggregates.** Phase-contrast images were used to segment microcolonies at exponential phase using MATLAB R2021b. The contrast was adjusted to the same range for each image by remapping the intensity to fixed values for each image (e.g. time points of a time-lapse). A Gaussian blur was applied to enhance the pixel detection during the binarization. To select the area of interest used for the segmentation and smooth the boundaries of the object, MATLAB's *strel* function was used to dilate and erode in a squared structuring element with a radius of 100 pixels. Each refined image was then used to detect the microcolonies, by first applying the Gaussian blur, adjusting the contrasts (MATLAB, *imadjust*), binarizing (*imbinarize*), filling holes (*imfill*), and smoothing the borders using the "strel" function with diamond structuring element with radius of 15 pixels. Objects of interest were then automatically detected using MATLAB's *regionprops* function. Unwanted objects and agarose crystals were manually removed. The biomass was defined as the pixel area of the segmented microcolony, projected in a 2D plane. To follow biomass growth over time, we selected only microcolonies with continuous presence in the field of view.

Stationary phase microcolonies and cell aggregates from liquid cultures were also segmented on images using MATLAB R2021b. Images were segmented using the eCherry fluorescence. The background intensity was determined by calculating the median value of the images in which the microcolonies and aggregates covered less than 50% of the field of view at each time point. The background intensity was then subtracted from the intensity measured at each pixel. Next, the MATLAB built-in function *im2bw* was used to create a binary mask, and to select the pixels corresponding to the biomass. To isolate the aggregates in liquid cultures, we filtered the segmented objects by pixel size and excluded objects with the size of a single bacteria from further processing.

Using the resulting masks and MATLAB's built-in function *regionprops*, we then extracted the fluorescence signal from each aggregate or microcolony (sum of the fluorescence intensity of all pixels divided by the surface area). To discriminate between the fluorescence signal of the bioreporters and autofluorescence from *P. putida* GB-1, we conducted control experiments with the untagged wild-type strain. The bioreporter fluorescence threshold was selected as the $99^{th}$ quantile of the fluorescence pixel intensity distribution of the wild-type, grown under the same condition as the bioreporter strains. All signal intensities are reported in arbitrary units (a.u.) (**Supplementary Fig. 20**).

**Image analysis of single cells from liquid cultures**. In the case of single (separated) cells on images, we used SuperSegger for segmentation[8], using a trained segmentation constant adapted to the size and shape of *P. putida* at 1000x magnification[9]. Mean cell fluorescence values (sum of the fluorescent pixels normalized by the cell area) were corrected by subtracting the median background signal outside the segmented cells. Wild-type *P. putida* GB-1 cell images in the eCherry channel were used to identify the cell auto-fluorescence (as the $99^{th}$ percentile of the wild-type fluorescence distribution) and the threshold above which we considered a 'true' eCherry bioreporter signal appearance. Segmented cells with mean eCherry fluorescence above the threshold were then considered as cells with active promoters (**Supplementary Fig. 21**). We further report the mean fluorescence bioreporter signal per reporting cell by averaging the individual cell fluorescence across all 'active' cells. The proportion of cells with active $P_{mnxG}$ or $P_{mcoA}$ promoters within a population is then the number of active cells divided by all segmented cells from the corresponding phase-contrast image. Representative results of the segmentation are shown in the false color figures (**Supplementary Fig. 21**).

**Quantification of Mn oxide precipitates on microcolonies by color analysis**. Color images were obtained in a separate experiment where more than 20 microcolonies were tracked. The surface area of microcolonies was measured using the same segmentation protocol described above. The proportion of cells within microcolonies that was covered in Mn oxide was quantified using color images obtained from the red, green, and blue color channels, using the 420 nm blue emission filter at 94% brightness, 510 nm green emission filter at 26% brightness, and 590 nm red emission filter at 69% brightness, respectively. All color channels were collected with 16 ms illumination time, 26 ms of camera exposure, and stored as 16-bit TIFF images. First, the images were inverted to obtain higher pixel intensities for the dark Mn oxide. The contrast was adjusted to the same scale by remapping the intensity to fixed values for each color channel. Images were then blurred, using a Gaussian blur, and the median value of the background of each color channel was subtracted. The brown precipitates of Mn oxide absorbed the transmitted light in the blue channel and did not show any absorption in the red channel (**Supplementary Fig. 22**). To separate the brown Mn oxide precipitates on the image from any color from the cells, we subtracted the blue channel intensity from the red channel, removed the noise from the resulting image using the *imnlmfilt* MATLAB function, and thresholded the results against the median intensity from images processed in the same manner at the time point prior to visible Mn oxide precipitation. The proportion of the microcolony covered in Mn oxides was then taken as the pixel sum of the identified precipitate 'area' using the regionprops function, divided by the total number of pixels identified as microcolony surface area. The detection limit of the Mn oxide, given the nanoscale dimensions of individual biogenic Mn oxide crystals (i.e., 6.25 ± 1.21 $nm^2$)[10], was estimated at the limited spatial resolution of the microscope (49,000 $nm^2$).

**Quantification of Mn oxide precipitates in liquid cultures by ICP-MS.** Aqueous Mn concentrations in the liquid cultures were measured by inductively coupled plasma mass spectrometry (ICP-MS, Agilent-7900). To discriminate between aqueous and solid phase Mn, we measured the aqueous and total Mn concentrations in sample aliquots. For aqueous Mn, we filtered 3 mL of the cell suspension using a 0.22 µm PES filter and acidified to 1 % nitric acid prior to analysis. For total Mn, 1 mL of the same cell suspension was digested by adding 30 µL of 65% nitric acid and 100 µl of 0.4 M oxalic acid, then filtered with a 0.22 PES filter to remove the biomass. The ICP-MS was equipped with a quartz spray chamber, a microMist concentric gas nebulizer, and nickel sampler and skimmer cones. ICP-MS analysis was performed in helium (He) mode, using a He flow rate of 4.5 mL $min^{-1}$ with 1.0 L $min^{-1}$ of argon carrier gas. The limit of quantification, which was calculated as 3.3 times the detection limit [11], was 0.04 µg $L^{-1}$ or 0.67 nM for Mn.

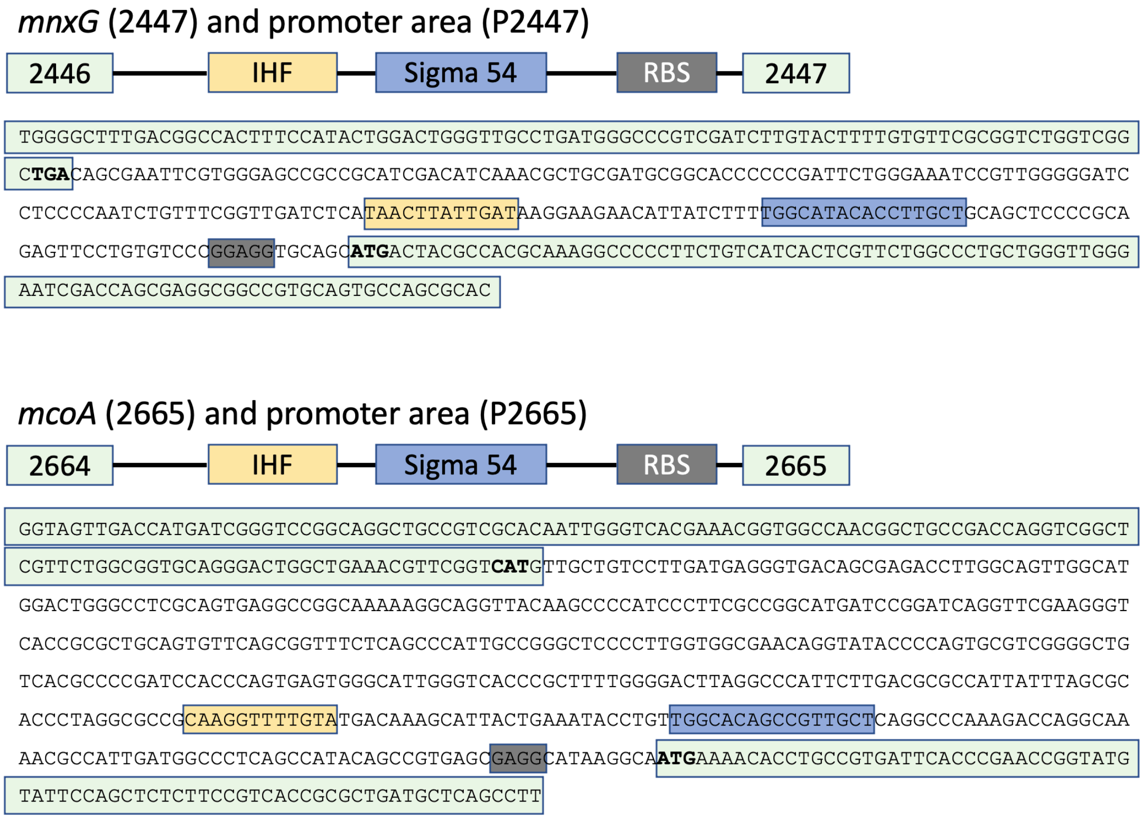


**Supplementary Fig. 1. Genetic sequences for *mnxG* and *mcoA*.** The top panel shows the sequence starting from gene 2446, with IHF, sigma 54 transcription factor, RBS, and 2447 (*mnxG*). The bottom panel shows the sequence starting from gene 2664, with IHF, sigma 54 transcription factor, RBS, and 2665 (*mcoA*).

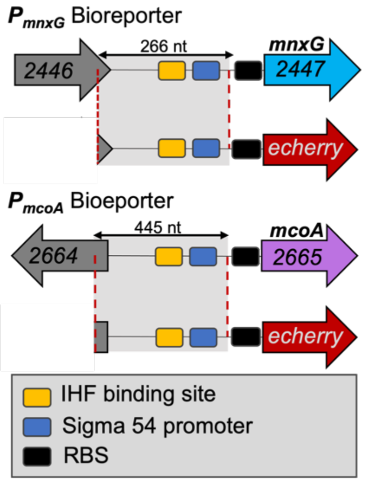


**Supplementary Fig. 2. Schematic representation of the single bioreporter systems (*$P_{mnxG}$* and *$P_{mcoA}$*) used to monitor manganese oxidase gene promoter activity in *Pseudomonas putida* GB-1.** Promoter regions (shown in grey and red dotted lines) devoid of RBS were fused to the *echerry* gene together with its RBS. Reporter fusions were inserted in a mini Tn5 delivery vector for subsequent single-copy integration into the genome of *P. putida* GB-1 containing the wild-type *mnxG* and *mcoA* loci. Genes are represented by arrow boxes. Each reporter was transformed into a separate strain. The scheme is not drawn to scale.

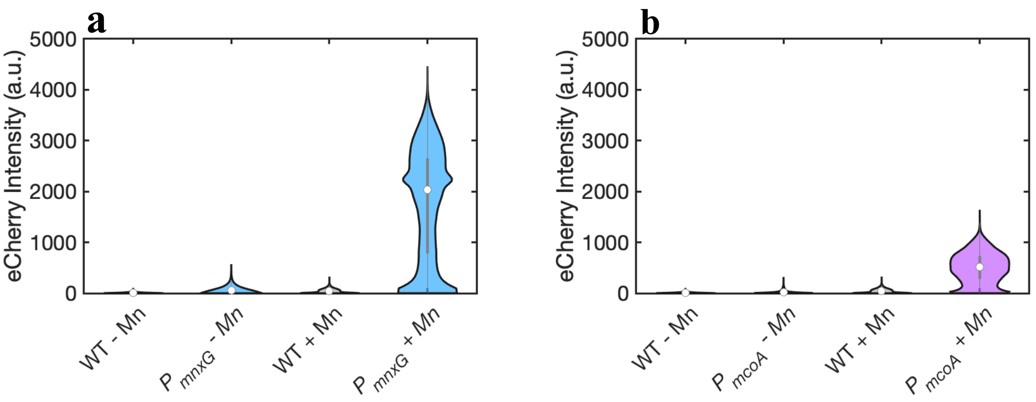


**Supplementary Fig. 3**. **Effect of Mn(II) on $P_{mnxG}$ and $P_{mcoA}$ activation in washed cultures**. Liquid cultures were washed at stationary phase and resuspended in MST salts with 50 µM $MnCl_2$ or without Mn. Samples were taken 24 h after washing for fluorescence measurements. (**a**) eCherry fluorescence signal distribution of wild type (gray) and bioreporter $P_{mnxG}$ (blue). (**b**). eCherry fluorescent signal distribution of wild type (gray) and $P_{mcoA}$ (purple). Data shown are for a single biological replicate for which more than 1000 cells were segmented.

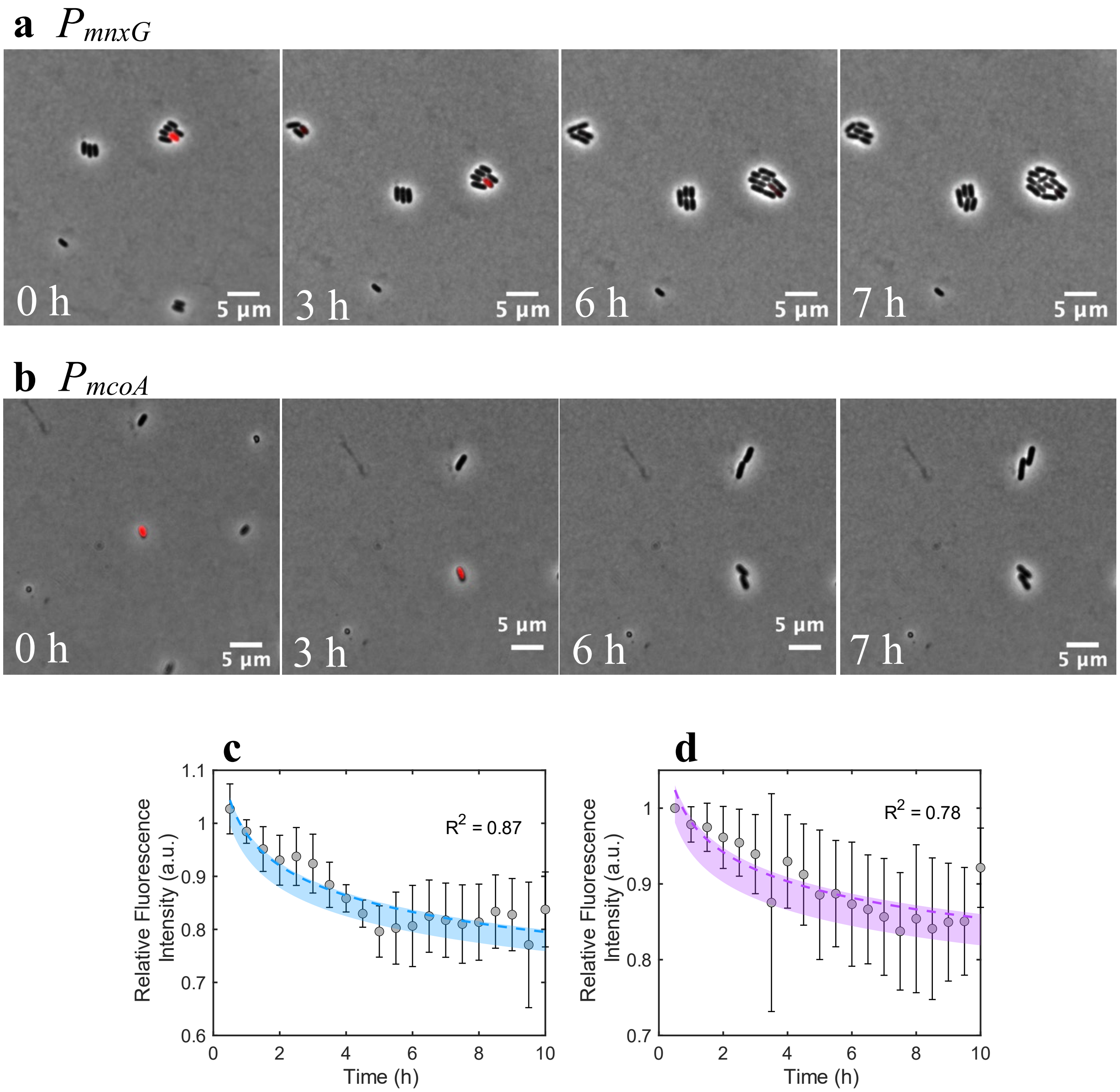


**Supplementary Fig. 4. Re-growth of $P_{mnxG}$ and $P_{mcoA}$ reporting cells on agarose patches.** Micrographs show the overlay of representative phase contrast and eCherry fluorescence images to illustrate the decrease in fluorescence intensity of $P_{mnxG}$ (**a**) and $P_{mcoA}$ (**b**) at 0 h, 3 h, 6 h, and 7 h. Reporting cells with active reporter $P_{mnxG}$ and $P_{mcoA}$ are shown in red. All images are rescaled to the same intensity. The relative average fluorescence intensity over time for three replicates of $P_{mnxG}$ (**c**) and $P_{mcoA}$ (**d**) are shown as grey symbols. The blue- and purple-colored areas show a decrease in average fluorescence signal with 95$^{th}$ confidence intervals, fitted as a power function. $R^2$ describes the power fitting relative to the average values. Results show a decrease in bioreporter activation after re-growth.

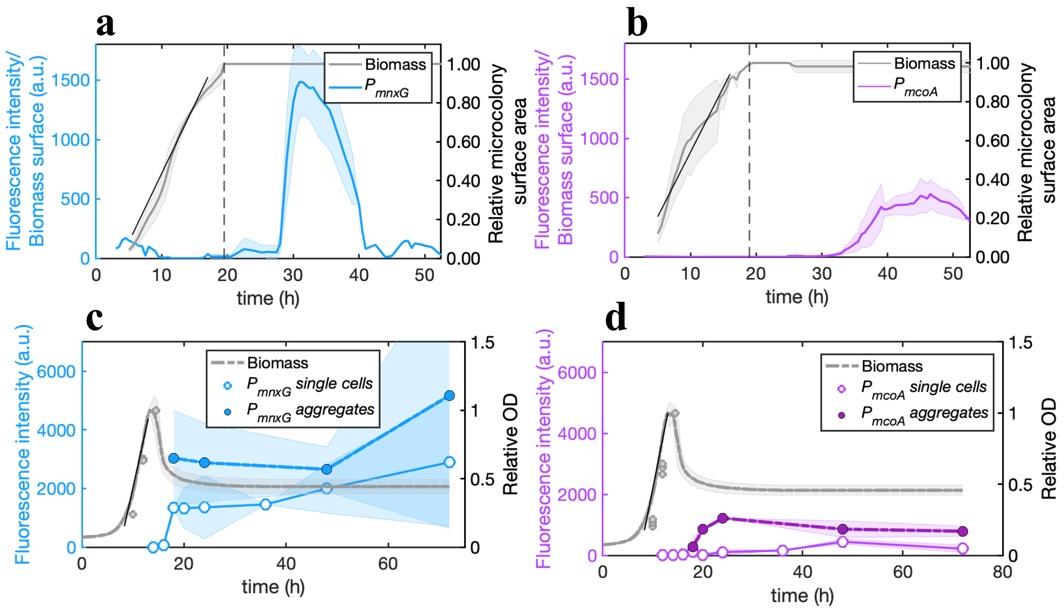


**Supplementary Fig. 5. Temporal activation of the Mn oxidase promoters of *P. putida* GB-1 in surface-grown microcolonies and liquid-suspended growth.** The sum of the fluorescence intensity normalized by microcolony size for (**a**) $P_{mnxG}$ and (**b**) $P_{mcoA}$ bioreporters, relative to the total surface area at entry into stationary phase at 19.5 h and 19 h, respectively (shown by the dotted vertical lines). The growth curve, calculated as the increase in microcolony surface, represents the average of 7 replicates for $P_{mnxG}$ and 21 replicates for $P_{mcoA}$. The fluorescence intensity of $P_{mnxG}$ (blue) is the average of 7 microcolonies, and $P_{mcoA}$ (purple) is the average of 21 microcolonies. Shaded areas represent the 95$^{th}$ confidence interval. Growth rates (μ) were calculated during exponential phase. The sum of the fluorescence intensity normalized by biomass for (**c**) $P_{mnxG}$ and **(d)** $P_{mcoA}$. The shaded area represents the standard deviation within triplicates. The plotted relative OD corresponds to the $OD_{600}$ normalized by the maximum $OD_{600}$. Growth curves for liquid cultures were measured in 5 replicates on a plate reader. The gray dots represent samples obtained from liquid flasks and compared to the growth observed in 96-well plates to assess for differences in growth in 96-well plates and Erlenmeyer flasks.

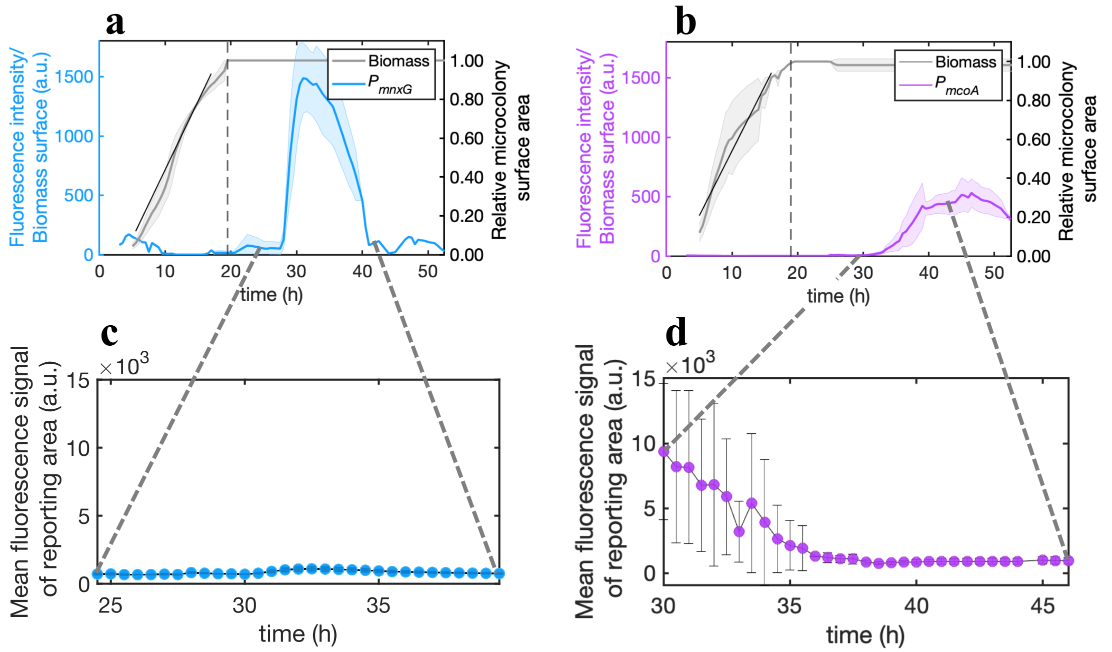


**Supplementary Fig. 6. Mean fluorescence per reporting microcolony area over time.** Overall reporter strain intensity $P_{mnxG}$ (**a**) $P_{mcoA}$ (**b**) relative to the total surface area at entry into stationary phase at 19.5 h and 19 h, respectively (shown by the dotted lines). Mean fluorescence signal per $P_{mnxG}$ (**c**) and $P_{mcoA}$ (**d**) reporting cell area. Results show no positive relationship between the average intensity with time, indicating that the overall increase in fluorescence intensity of $P_{mnxG}$ and $P_{mcoA}$, observed in the **Main text, Figure 2**, is due to an increase in the proportion of the population activating the promoters $P_{mnxG}$ and $P_{mcoA}$, and not due to an increase in fluorescence intensity per reporting cell. Note: the mean fluorescence signal of $P_{mcoA}$ reporting cells from 30 h to 33 h comes from less than 0.7% of the population since most of the microcolony has inactive $P_{mcoA}$.

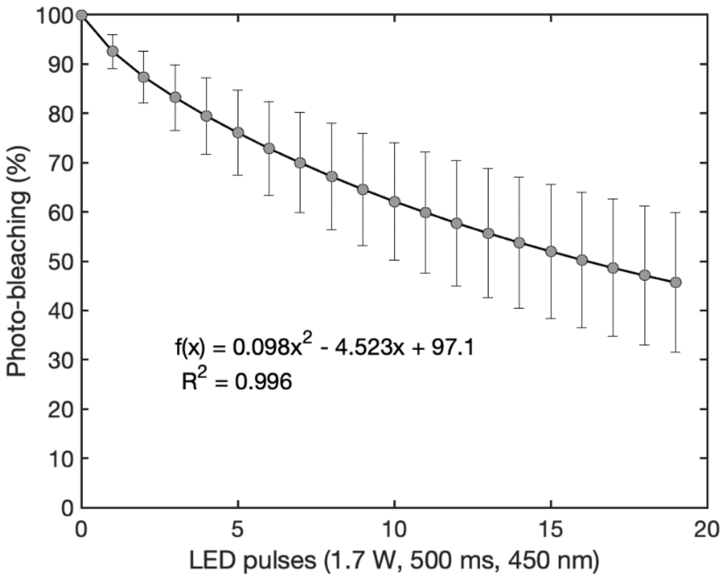


**Supplementary Fig. 7**. **Photobleaching due to prolonged LED exposure.** Photobleaching of 4 different sizes aggregates of $P_{mnxG}$. Aggregates were exposed to pulses of 500 ms LED in eCherry channel (emission at 450 nm) at 50% laser intensity (1.7W). The inset equation shows the second-degree fit for the average bleaching. The photobleaching rates measured on 72 h old microcolonies are calculated as 3.70 %/h, on average, and matched the decrease in $P_{mnxG}$ intensity calculated from 31.5 h to 35.5 h (**Fig. 2a**). After this 4 h time window, the fluorescence intensity decreased rapidly at a rate of 16.34 %/h, until the fluorescence intensity was no longer detectable (**Fig. 2a**), showing inactivation of the promoter. The fluorescence from $P_{mcoA}$, after 47 h on solid surface, decreased at a rate of 6.78 %/h (**Fig. 2b**), which was greater than the experimentally determined photobleaching rate, suggesting that $P_{mcoA}$ activation also ceased.

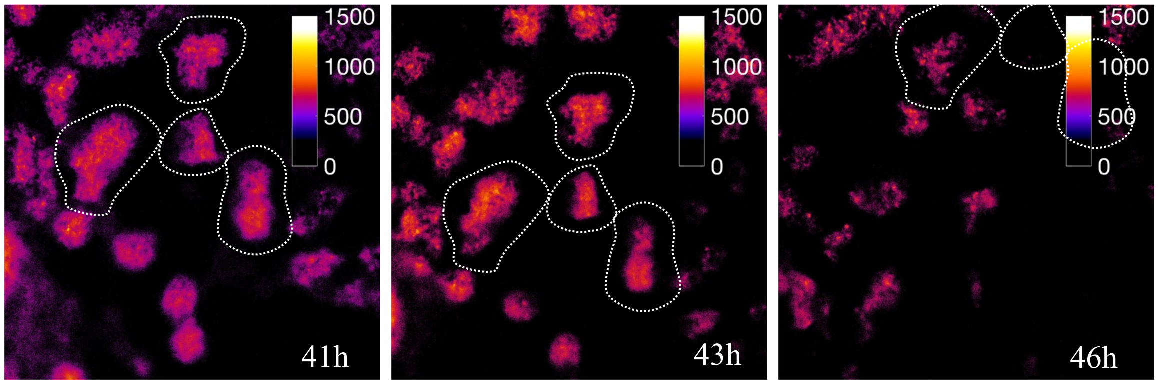


**Supplementary Fig. 8. $P_{mcoA}$ activation pattern from steady state to photobleaching.** The proportion of the microcolonies with active $P_{mcoA}$ does not increase further after reaching steady state (ca. 41 h) but decreases over time due to photobleaching. The shift in the location of the microcolonies within the frame is due to the drying of the agarose patch over time during extended time-lapses, shrinking it and shifting it from its set position.

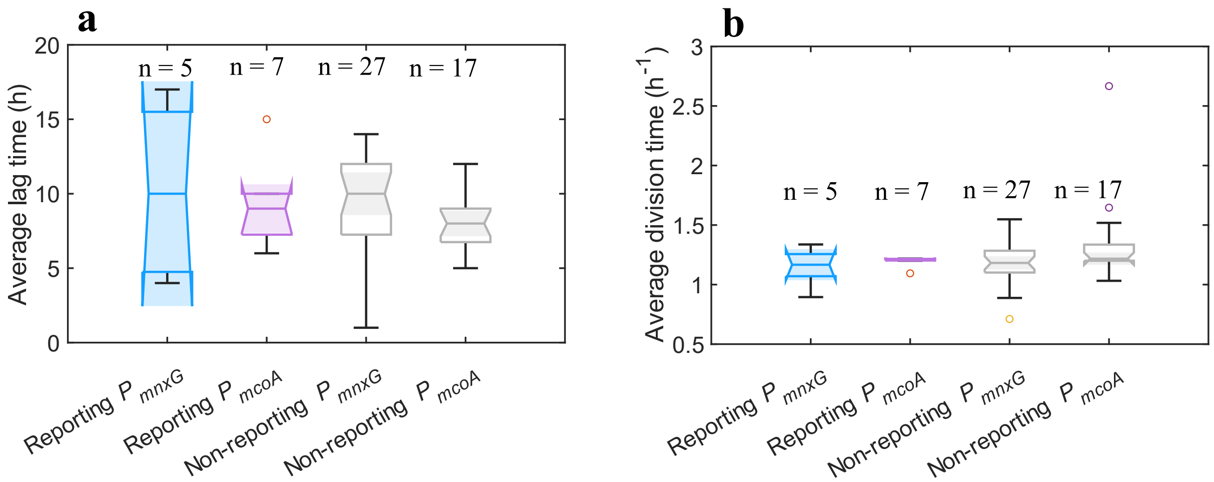


**Supplementary Fig. 9. Viability of reporting and non-reporting cells.** Cells were grown in MSTA containing 50 μM $MnCl_2$ for 48 h to generate a subpopulation of reporting cells. These cells were then washed and transferred to a 1% agarose patch containing 5 mM L-Arginine in MSTA. Cell division was monitored over 24h with a measurement made every 10 min and used to determine the average lag phase (**a**) and average division time (**b**) for reporting and non-reporting cells. ANOVA showed no significant differences between the reporting and non-reporting cells for both lag phase length ($F>0.282$, df=3) and division time ($F>0.099$, df=3). Note that the non-reporting cells may be expressing the untagged genes. However, the uniform distributions of the bioreporters suggest that the untagged genes did not skew the distribution.

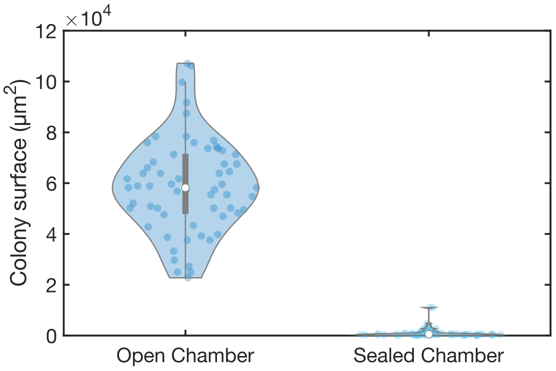


**Supplementary Fig. 10. Surface area of the colonies** cultured on MSTA solid surfaces (1% agarose) with 50 μM $MnCl_2$ in open and closed chambers. Data shows a greater than 100-fold increase in the mean surface areas of the microcolonies grown open to the air compared to the ones grown in closed chambers, which indicates that oxygen is the growth limiting factor in the closed microscope chambers.

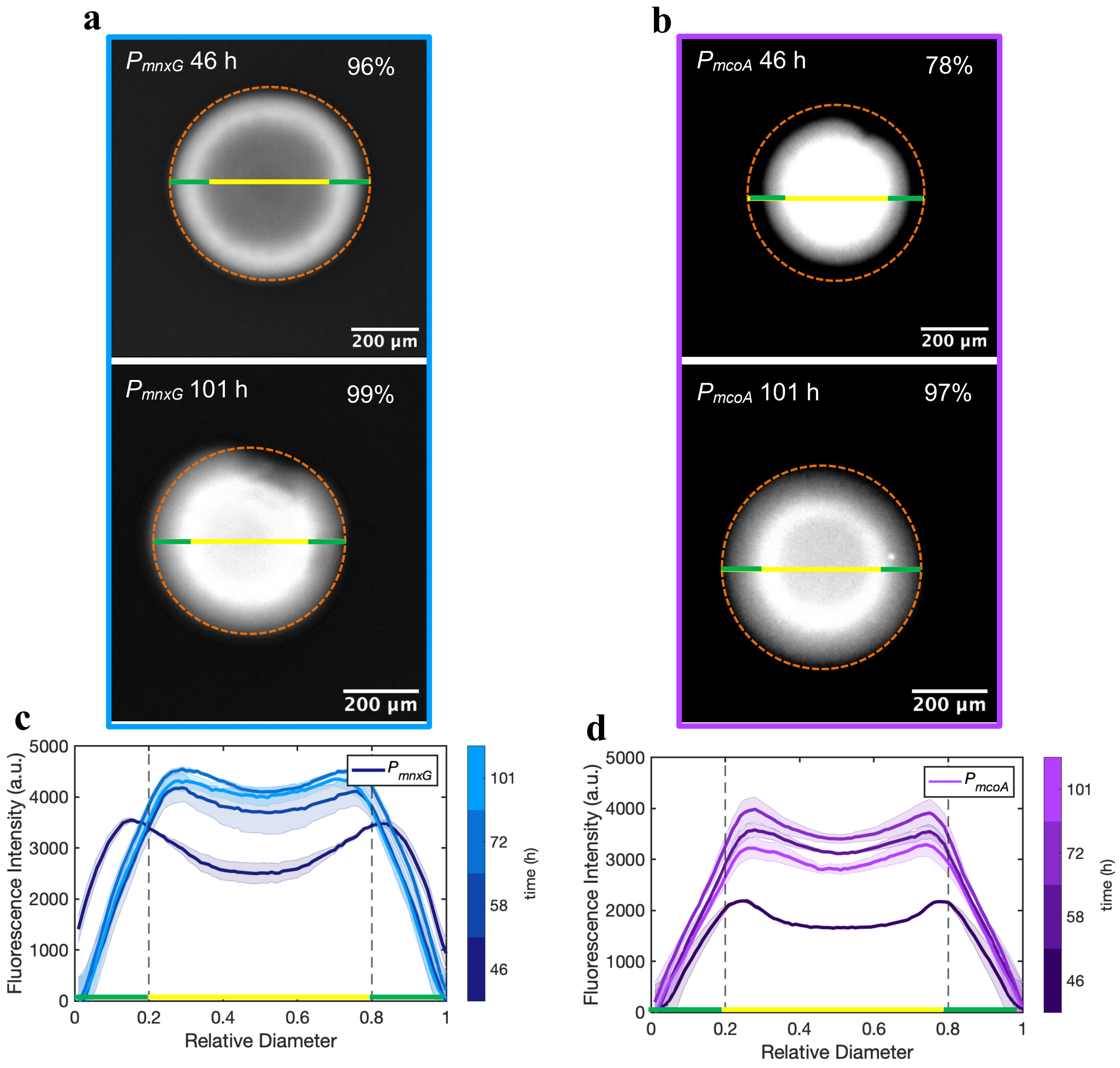


**Supplementary Fig. 11. $P_{mnxG}$ and $P_{mcoA}$ activation patterns in open chambers colonies.** Activation pattern of $P_{mnxG}$ at 46 h and 101 h (**a**). Activation pattern **of $P_{mcoA}$ at** 46 h and 101 h (**b**). The dotted orange lines represent the boundaries of the microcolony. The percentages represent the percent coverage of the microcolony with active bioreporter. Intensity profiles along the diameter of the microcolonies are shown in panel (**c**) for $P_{mnxG}$ and panel (**d**) for $P_{mcoA}$. The profiles are the average of 5 replicates for each strain and each time point. The center of the profile is represented in red, and the outer rim is represented in green. Results show that $P_{mnxG}$ activates first in the outer rim and then fills in. $P_{mcoA}$ activates first in the center and then slowly progresses outwards. These patterns are consistent with the ones observed in the sealed-chambers microcolonies (**Main text, Fig. 3.**) and confirm that the fluorescence patterns are induced by the microcolony and not an artifact of the restrictive conditions found in sealed chambers.

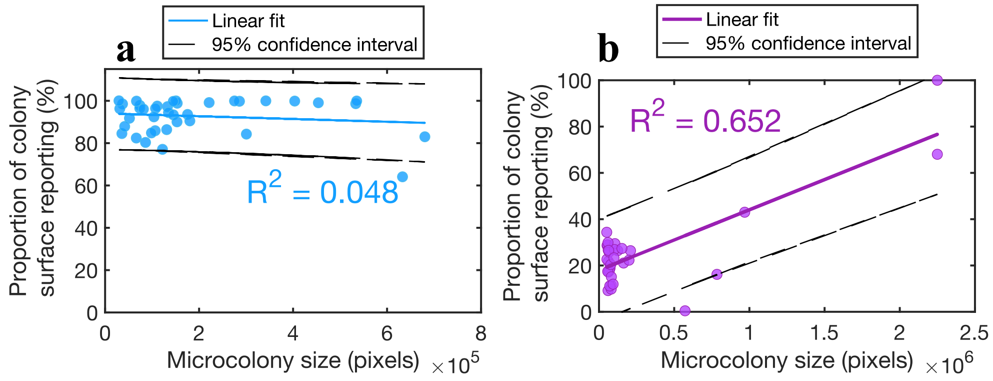


**Supplementary Fig. 12. Relationship between colony size and proportion of the microcolony surface reporting.** $P_{mnxG}$ strain shows no linear relationship with colony size (n = 39) (**a**), whereas $P_{mcoA}$ strain shows a linear relationship with colony size (n=33) with a p-value < 0.01 (**b**). Data originate from the 48 h time points in the time-lapse experiments presented in **Figs. 2** and **4** (Main text), with most of the $P_{mcoA}$ size variability coming from the time-lapse in **Fig. 2**.

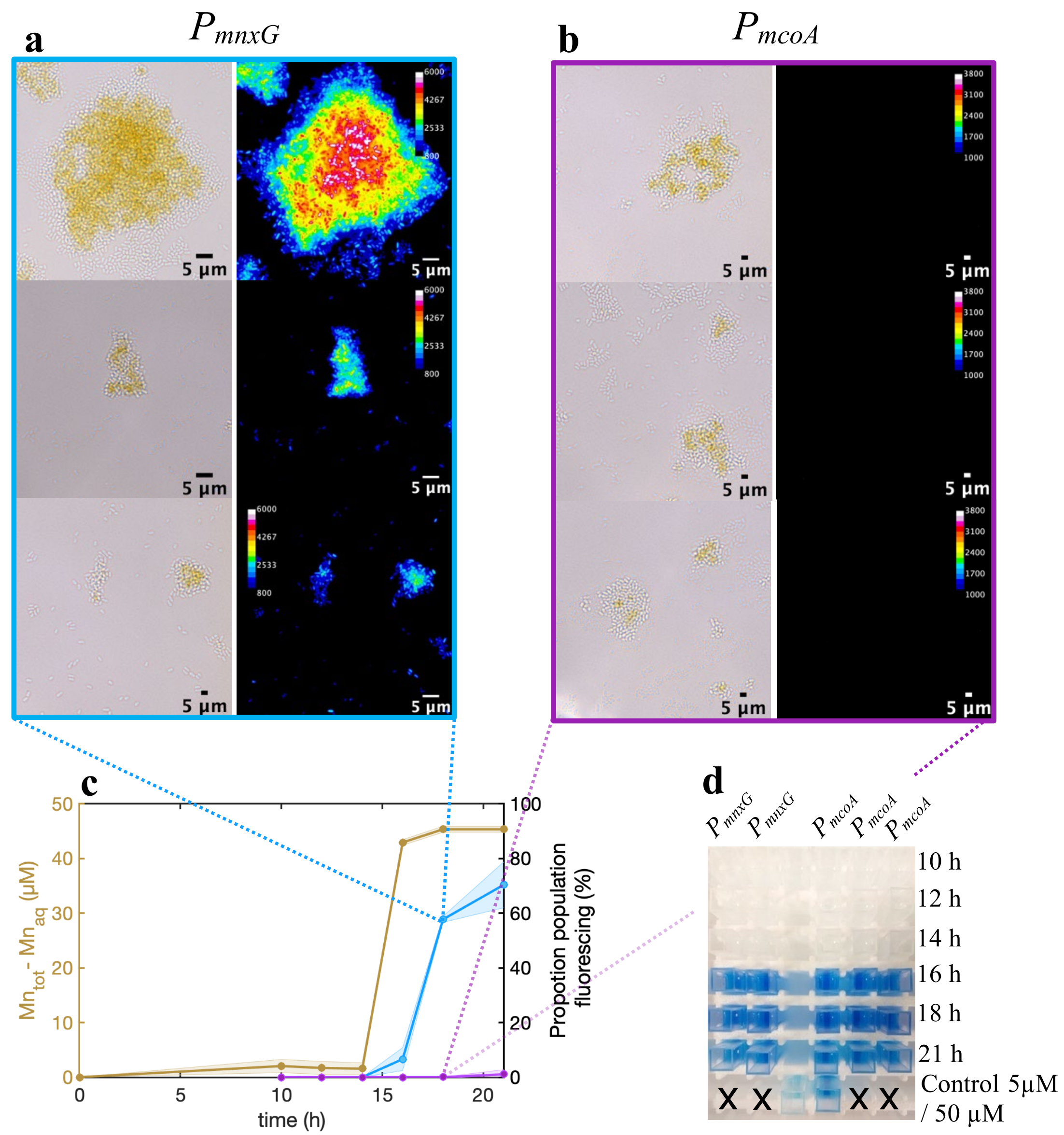


**Supplementary Fig. 13. Mn(II) sorption and proportion of the population at mid-exponential phase and early stationary phase.** (**a**) Representative brightfield color images of single cells and aggregates ($P_{mnxG}$), showing Mn oxide precipitates accumulating in the center of the aggregates (left column) and the corresponding $P_{mnxG}$ reporter fluorescence (right column) at 18 h. (**b**) Representative brightfield color images of single cells and aggregates ($P_{mcoA}$), showing Mn oxide precipitates accumulating in the center of the aggregates (left column) and the corresponding $P_{mcoA}$ reporter fluorescence (right column) at 18 h. (**c**) Proportion of population fluorescing over time for $P_{mnxG}$ and $P_{mcoA}$. Patterns observed in **a** and **b**. indicate that the initial Mn oxide precipitation is attributed to $P_{mnxG}$ and MnxG since no fluorescence was observed for PmcoA, but Mn oxides were present. The amount of Mn in the aqueous phase shows at most 3.9 ± 2.2% of sorption to the biomass prior to gene activation. (**d**) LBB colorimetric assay shows Mn oxide production starts at 16 h and coincides with $P_{mnxG}$ activation. The LBB controls used δ-$MnO_2$ in MST salts at 5 µM and 50 µM concentrations. Data is shown in duplicates for $P_{mnxG}$ and triplicates for $P_{mcoA}$.

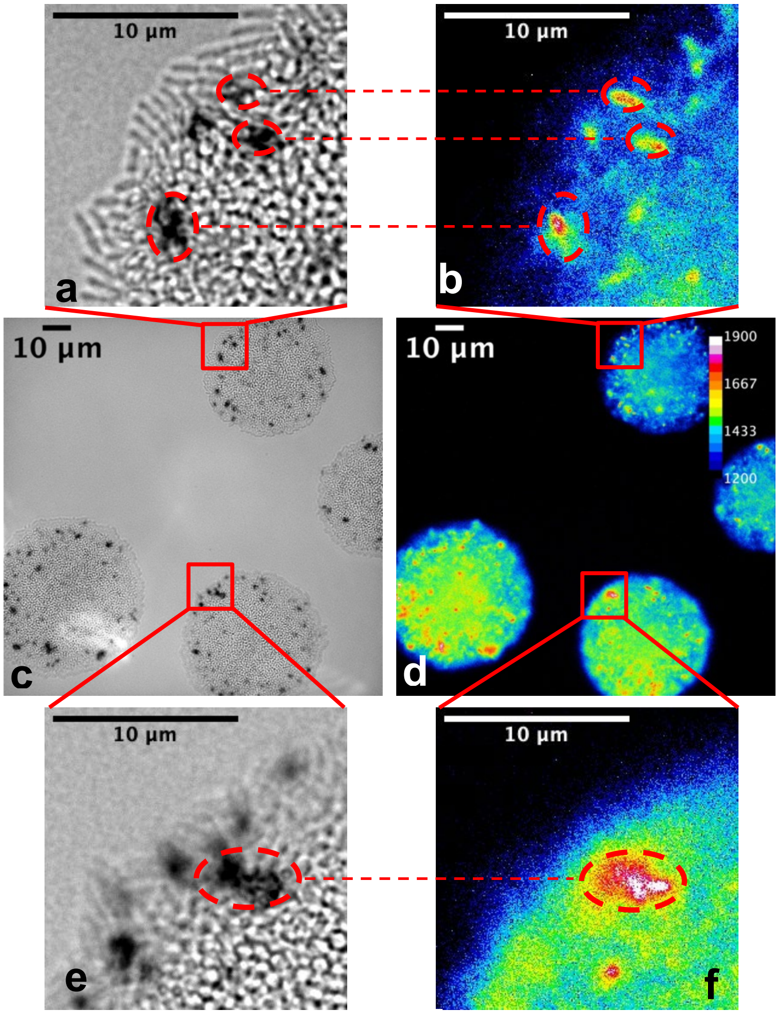


**Supplementary Fig. 14. Micrographs showing co-location of hotspots of Mn oxide precipitates and promoter activation for the $P_{mnxG}$ bioreporter.** Brightfield images of microcolonies acquired at 40 h in the blue channel to delineate the Mn oxides (**c**) and heatmap of the fluorescence signal at 37 h (**d**). Panels **a**-**b** and **e**-**f** provide a close-up view of hotspots of Mn oxide (**a**, **e**) and $P_{mnxG}$ fluorescence (**b**, **f**).

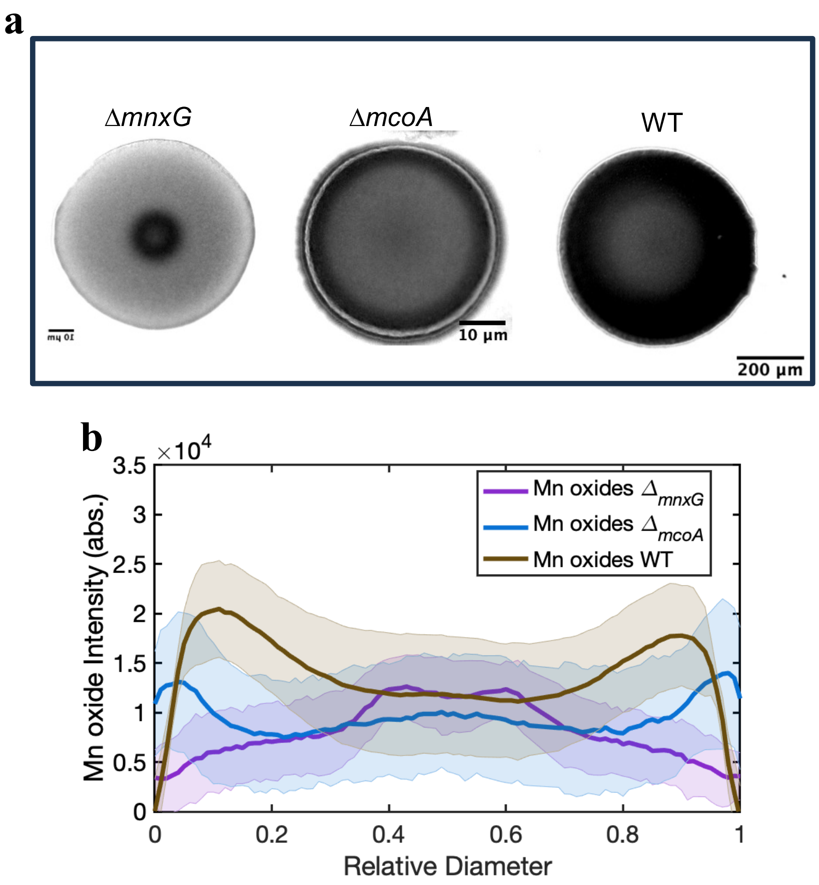


**Supplementary Fig. 15. Mn oxide precipitation by the single knockouts and the wild type of *Pseudomonas putida* GB-1.** The red subtracted blue channels or the Δ*mnxG,* Δ*mcoA*, and WT (**a**), highlighting the location of the Mn oxide precipitates. Average intensity profiles of the inverted color images for the single knockouts and WT (**b**). Averages of triplicates and the standard deviation are shown in a shaded area. Results show that McoA (Δ*mnxG*) produces Mn oxides only in the center, whereas MnxG (Δ*mcoA*) precipitates the oxides in the outer rim. The wild type shows Mn oxide precipitates in the whole colony, suggesting that each Mn oxidase is specific for local chemical conditions. The images were acquired at steady state (101 h). Note: the lighter shade in the center for the WT colony is an artifact of the image processing, resulting from darker biomass and thick layer of Mn oxides in the center that also shows a signal in the blue channel.

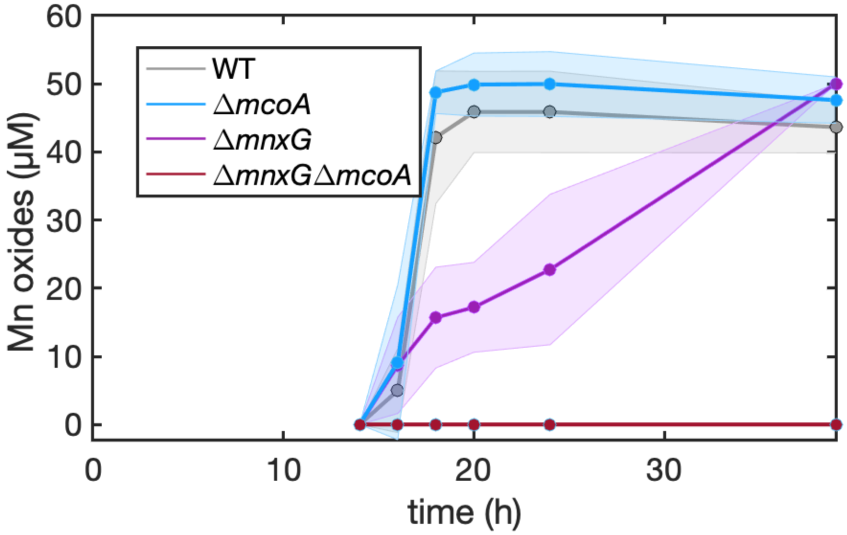


**Supplementary Fig. 16. Mn oxidation kinetics of the wild type and single and double gene knockout strains.** Data show that the Mn oxidation kinetics of the wild-type strain matches the Mn oxidation kinetics for the strain containing MnxG alone. While the strain containing McoA can eventually remove the 50 μM Mn(II) from the solution, the kinetics are much slower. The Mn oxide precipitated (y-axis) was calculated as the difference between the acid-digested Mn(II) and the aqueous Mn(II), and measured by ICP-MS. Shaded areas show the standard deviation within 3 replicates.

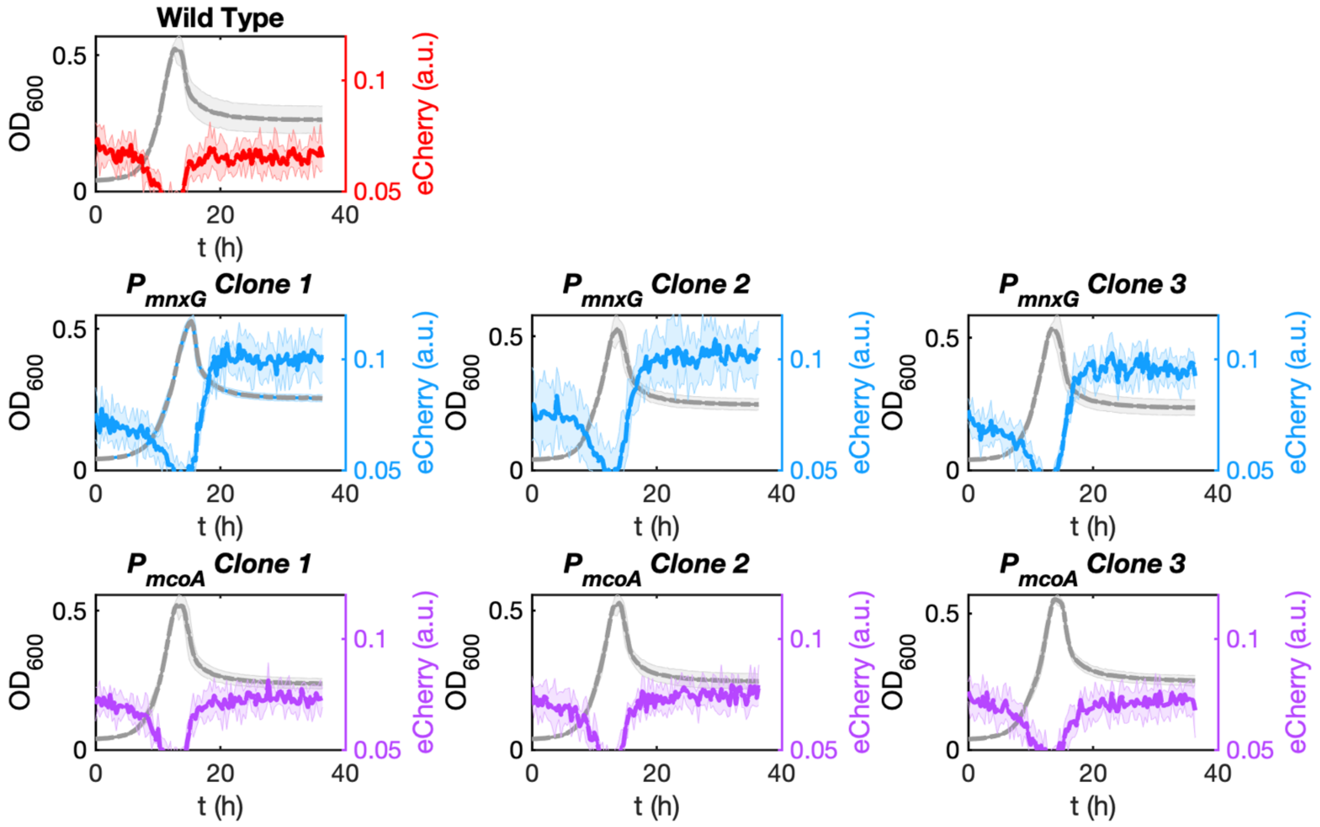


**Supplementary Fig. 17. Growth and fluorescence of the-wild type (WT) and the bioreporter clones.** Growth curves and fluorescence measurements were obtained using Varioskan LUX Multimode Microplate Reader at 30°C and under constant orbital shaking (160 rpm). Six replicates were measured for each strain in MSTA supplemented with 50 µM $MnCl_2$. Stationary phase was reached at 13h for all strains and replicates apart from the $P_{mnxG}$ clone 1, reached stationary phase at 15h.

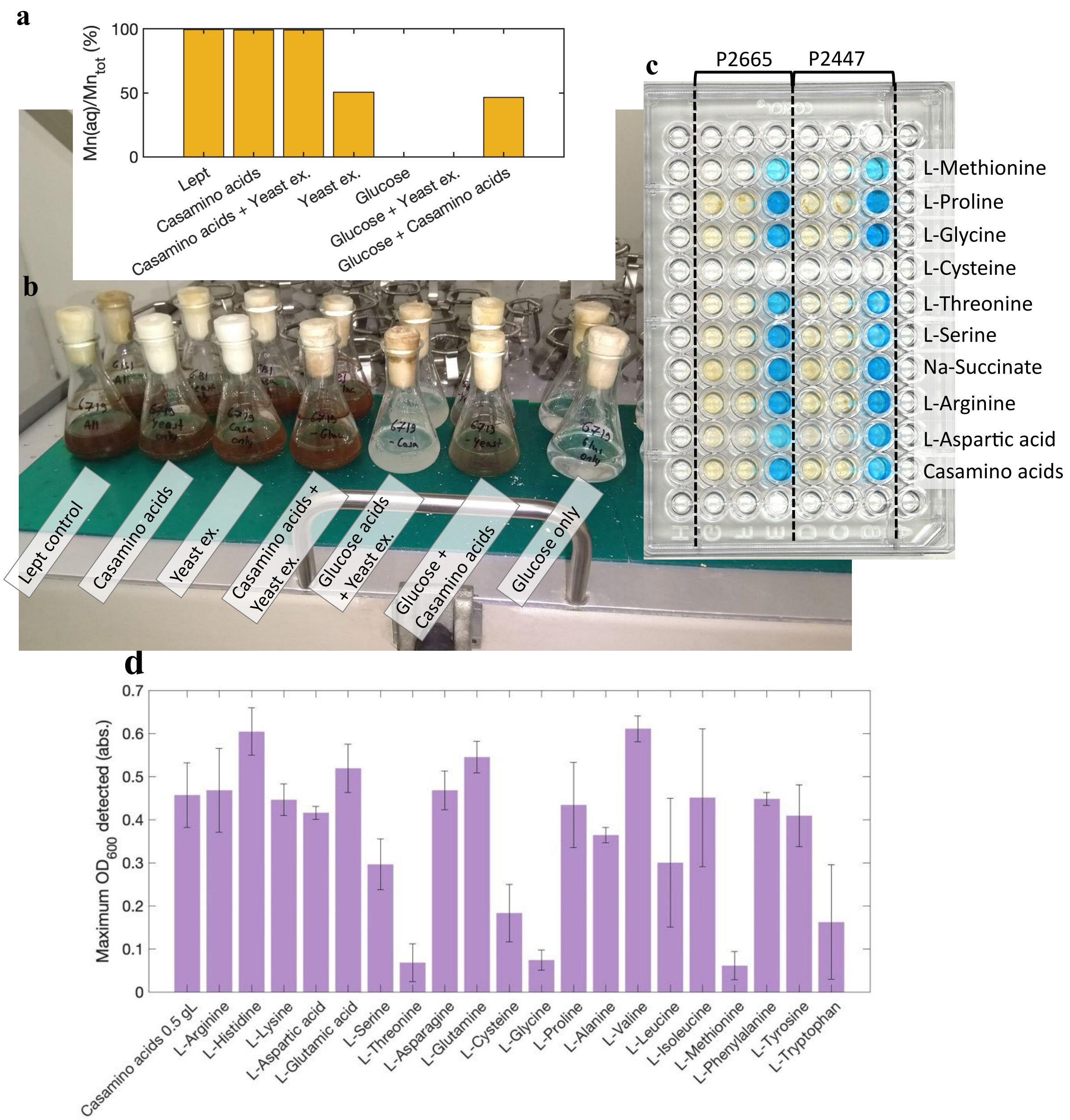


**Supplementary Fig. 18. Development of MSTA growth medium.** (**a**) Quantification of Mn oxides produced in Leptothrix salt supplemented with either Casamino acids, Glucose, or Yeast extract, or a mix. The percent of Mn oxides was calculated by the difference between total Mn and aqueous Mn, normalized to the total Mn. Quantification for a single replicate. (**b**) Visual brown Mn oxide precipitates in Leptothrix salts containing Casamino acids, Glucose, Yeast extracts, or a mix. Quantification in 6 replicates with LBB assay in duplicates. Results indicate that Casamino acids are the main inductors of Mn oxidation, whereas Glucose prevented Mn oxide precipitation. (**c**) Single amino acids (5 mM) were tested in MST salts and incubated in the plate reader at 30° and 180 rpm for 48 h, in triplicates per bioreporter strain. The amounts of Mn oxide precipitates were qualitatively assessed using LBB colorimetric assay. (**d**) Maximum growth was measured for the tested amino acids (each amino acid concentration was 5 mM). Results show that L-Arginine is the preferred carbon source with strong Mn oxide precipitation. All tested media were buffered at pH 7.0 using HEPES.

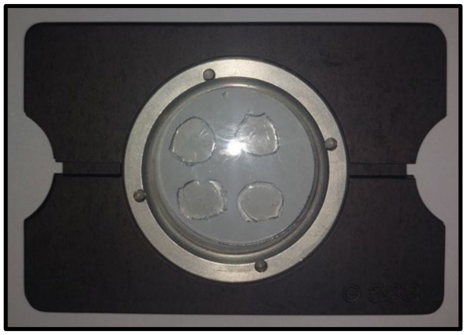

**Supplementary Fig. 19. Microscopy chamber** (Helmut Saur Laborbedarf, Germany), containing four agarose patches. The patches contained 1% agarose and MSTA growth medium with 50 µM $MnCl_2$. Two needles were introduced through open slits on the side of the chamber to improve oxygenation.

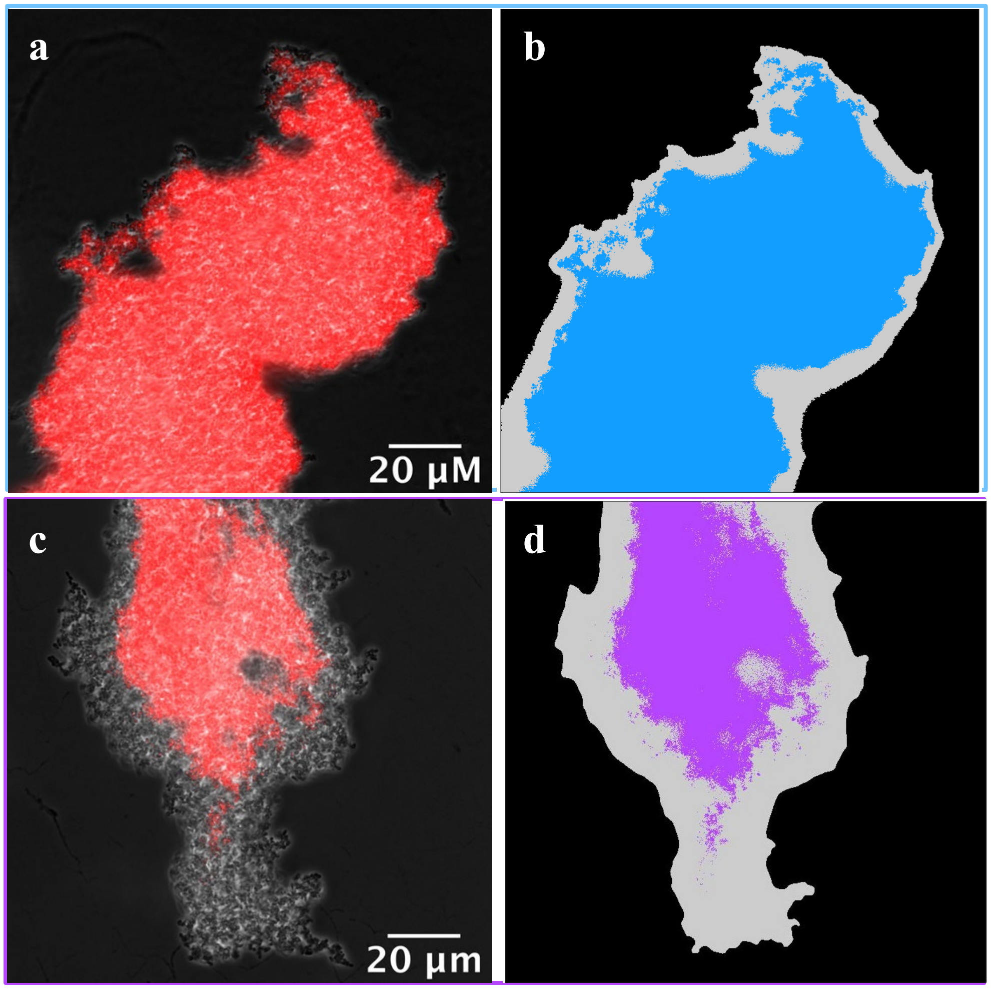


**Supplementary Fig. 20. Analysis of the surface covered by reporting cells in aggregates.** Representative image of phase contrast overlapped with eCherry fluorescence (in false red color, after applying the threshold) of the strain $P_{mnxG}$ (**a**) grown in liquid with 50 μM $MnCl_2$ and the corresponding segmented mask (**b**). Representative image of phase contrast overlapped with eCherry fluorescence of the strain $P_{mcoA}$ (**c**) grown in the same conditions and its corresponding segmented mask (**d**). The gray shows the aggregate area and the active reporters in color. The biomass segmentation is less accurate at the edges, with more biomass segmented than the reality. This results in a slight underestimation of the proportion of the biomass showing $P_{mnxG}$ or $P_{mcoA}$ activity.

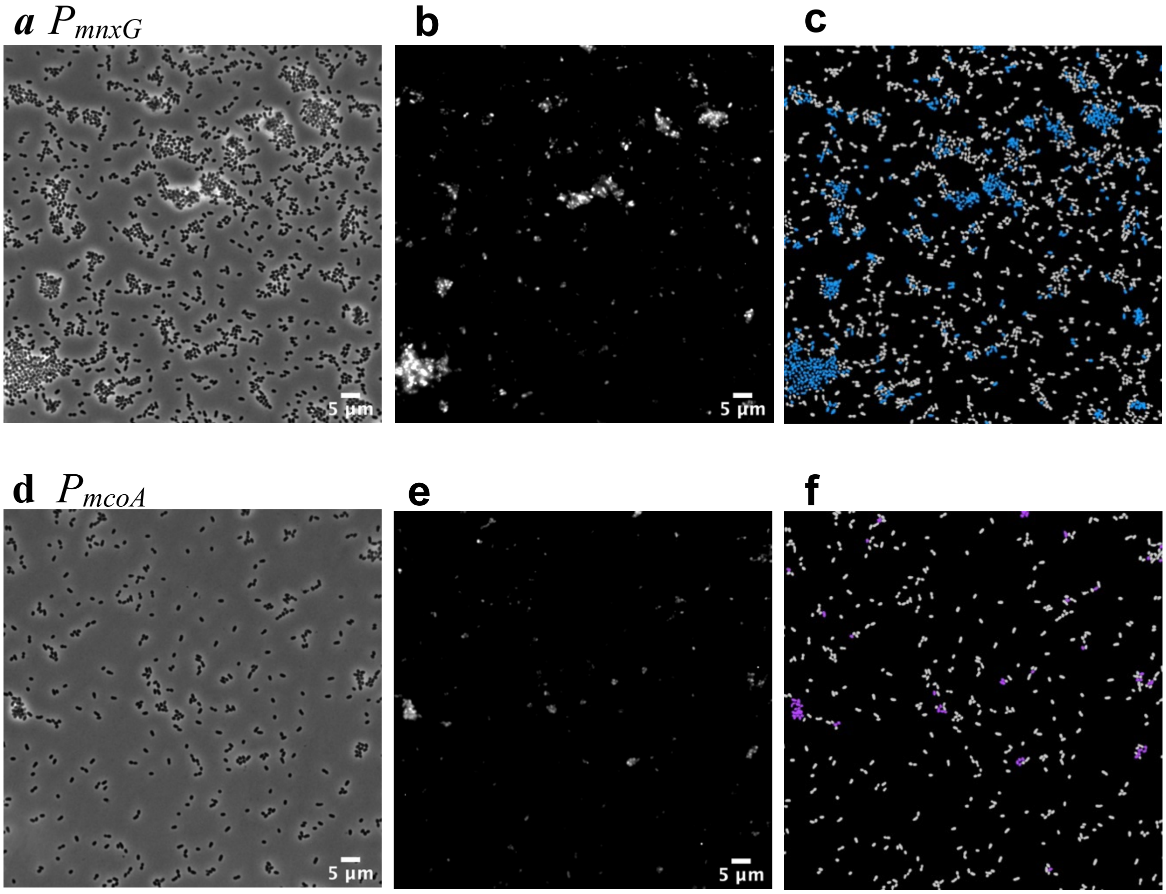


**Supplementary Fig. 21. Representative image segmentation of single cells.** (**a**) $P_{mnxG}$ phase contrast image. (**b**) $P_{mnxG}$ eCherry fluorescence image. (**c**) $P_{mnxG}$ false-colored image output of the segmentation and identification of fluorescing cells, with 28.2% of the population fluorescing (in blue). (**d**) $P_{mcoA}$ phase contrast image. (**e**) $P_{mcoA}$ eCherry fluorescence image. (**f**) $P_{mcoA}$ false-colored image output of the segmentation and identification of fluorescing cells, with 10.5% of the population fluorescing (in purple). All images acquired at 1000x, in the MSTA containing 50 µM Mn(**II**).

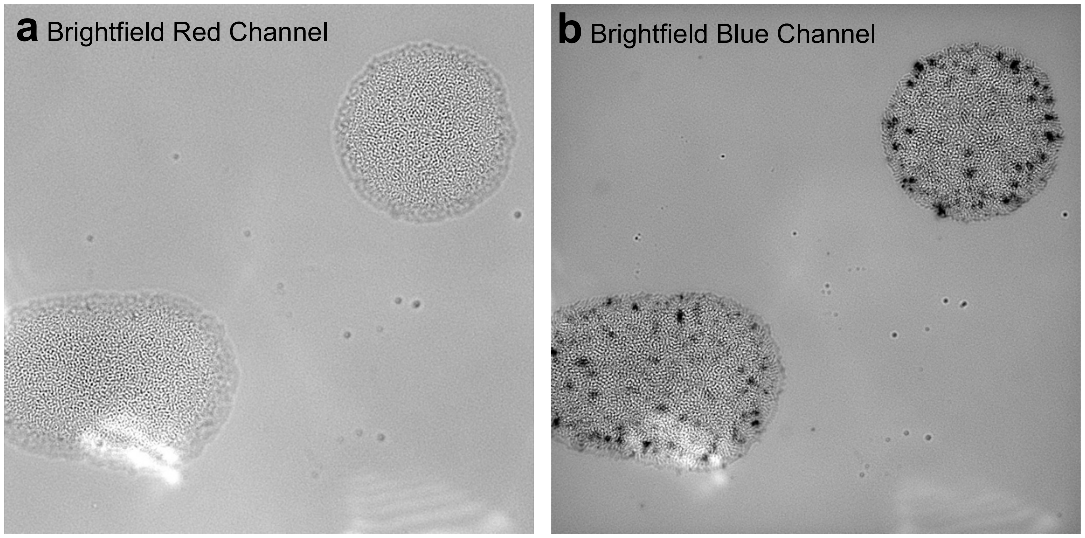


**Supplementary Fig. 22. Color channel comparison for Mn oxide quantification.** (**a**) Brightfield image acquired in the red channel. (**b**) Brightfield image acquired in the blue channel. The dark shadows indicate the location of the Mn oxide precipitates. Both channels are from the same original RGB color image.

**Supplementary Table 1. Strains used in this study.**

| Strain Label | Description | Use |
|---|---|---|
| WT | *Pseudomonas putida* GB-1 Wild-type | Autofluorescence control |
| $P_{mnxG}$ | *P. putida* GB-1::P2447_eCherry | Follow activation of promoter $P_{mnxG}$ (*mnxG*) |
| $P_{mcoA}$ | *P. putida* GB-1::P2665_eCherry | Follow activation of promoter $P_{mcoA}$ (*mcoA*) |
| *ΔmnxG* | *P. putida* GB-1 single knockout of *mnxG* | Follow Mn oxidation by McoA |
| *ΔmcoA* | *P. putida* GB-1 single knockout of *mcoA* | Follow Mn oxidation by MnxG |
| *ΔmnxGΔ mcoA* | *P. putida* GB-1 double knockout of *mnxG* and *mcoA* | Mutant without the capacity to oxidize Mn |
| pBAM1 | Host plasmid containing the PmnxG_eCherry or PmcoA_eCherry cassettes, and Tn5 transposon. Stored in *E. coli* DH5α | Construction of the fusion reporter strains |

**Supplementary Table 2. Primer used to construct the bioreporters and the knockout strains**

| Bioreporter strains | InFusion reaction to insert the promoter and *eCherry* gene into the pBAM1 vector digest with SmaI |
|---|---|
| PputGB1_2447.F | TTCGAGGCATGCCTGCAGCCCGGCCACTTTCCATACTGGACT |
| PputGB1_2447.R | ACCATGGCAGGTGCTCCTTCT CCGGGACACAGGAACTCT |
| PputGB1_2665L.F | TTCGAGGCATGCCTGCAGCCC CAGGCTGCCGTCGCACAAT |
| PputGB1_2665.R | ACCATGGCAGGTGCTCCTTCT GCTCACGGCTGTATGGCTGA |
| eCherry.F | agaa**ggag**cacctgccatggt |
| eCherryTn5.R | AATCAGAATTCGAGCTCGCCC ttatttgtacagctcatccatgcca |

| Deletions (Knockouts) | Cloning of upstream and downstream homologous fragment into pJP5603-ISceI |
|---|---|
| D2447UP.for | AGGGATAACAGGGTAATCTGAATTCGCGCTTGACCCCACCCCTGAA[2] |
| D2447UP.rev | TTTCCGTCACTGCGCCGGGGC GCGTGGCGTAGTCATGCTGCA |
| D2447DW.for | GCCCCGGCGCAG**TGA**CGGAAA |
| D2447DW.rev | GAAGCTTGCATGCCTGCAGGTCGAC GACCGTGTGGCGCAGCTGAT |
| D2665UP.for | AGGGATAACAGGGTAATCTGAATTC TCGAAGAAGGCCAGCTCCAA[2] |
| D2665UP.rev | ACGCGTCGGTTACTTGATGCTGAT GGCAGGTGTTTTCATTGCCTT |
| D2665DW.for | ATCAGCATCAAG**TAA**CCGACGCGT |
| D2665DW.rev | GAAGCTTGCATGCCTGCAGGTCGACCGGAAAATACGACAGTAAGAT |

**Supplementary Table 3. Defined growth medium composition of Mineral Salts and Traces L-Arginine (MSTA).** All solutions were prepared with Milli-Q ultrapure water (18.2 MΩ·cm), and filter-sterilized with a 0.2 μm PES filter membrane. All reagents were stored at 4°C and prepared with ACS-grade chemicals. Final pH 7.0.

| **Major Elements** | **Concentration in the growth medium** |
|---|---|
| $CaCl_2 \cdot 2H_2O$ | 0.4 mM |
| $MgSO_4 \cdot H_2O$ | 0.25 mM |
| $Na_2HPO_4$ | 0.25 mM |
| $KH_2PO_4$ | 0.15 mM |
| 1:2 Fe(III)-EDTA | prepared by reacting 20 μM $FeCl_3 \cdot 6H_2O$ with 40 μM EDTA, adjusted at pH 6.5 with NaOH |
| HEPES buffer | 10 mM, prepared by adjusting the pH to 7.0 with NaOH |
| $(NH_4)_2SO_4$ | 5 mM |
| **Trace Elements** | |
| $CuSO_4 \cdot 5H_2O$ | 40 nM |
| $ZnSO_4 \cdot 7H_2O$ | 273 nM |
| $CoCl_2 \cdot 6H_2O$ | 84 nM |
| $NaMoO_4 \cdot 2H_2O$ | 53.7 nM |
| **Carbon Source** | |
| L-Arginine* | 0.5 mM |

*L-Arginine was omitted for the preparation of the MSTA salts solution.

**Supplementary Table 4. Characteristics of the bioreporter strains in MSTA (*SI Material and Methods*) containing 50 µM $MnCl_2$.** Growth rates (calculated from the optical density at the exponential phase) and fluorescence were recorded using a Varioskan LUX Multimode Microplate Reader in 96-well plates. Mn oxide precipitation rates, onsets, and time to full oxidation were measured in batch experiments (Erlenmeyer flasks) as described in the method section.

| | Growth rates ($h^{-1}$) | Fluorescence intensity (a.u.) | Mn oxide precipitation rate (µM/h) | Mn oxidation onset (h) | Time to full oxidation(h) |
|---|---|---|---|---|---|
| WT | 0.33 | - | 13.6 ± 5.7 | 16 ± 0.0 | 20.5 ± 2.2 |
| $P_{mnxG}$ - Clone 1 | 0.26 | 2.45 | 13.2 ± 8.1 | 16.5 ± 1.7 | 20 ± 2.8 |
| $P_{mnxG}$ - Clone 2 | 0.33 | 2.31 | 11.0 | 12 | 18 |
| $P_{mnxG}$ - Clone 3 | 0.33 | 2.39 | 22.6 | 16 | 18 |
| $P_{mcoA}$ - Clone 1 | 0.32 | 2.09 | 12.6 ± 5.5 | 16 ± 1.4 | 21 ± 3.0 |
| $P_{mcoA}$ - Clone 2 | 0.31 | 1.87 | 12.9 | 14 | 18 |
| $P_{mcoA}$ - Clone 3 | 0.28 | 1.78 | 23.9 | 16 | 18 |
